# Linker-Length Landscape Mapping Enables Coupling of Diverse Synthetic Chemically Induced Dimerization Systems to Molecular Readouts

**DOI:** 10.64898/2026.07.01.735888

**Authors:** Yuxin Pan, Shoukai Kang, Davi Nakajima An, Yixuan Yu, Frank DiMaio, Liangcai Gu

**Affiliations:** Department of Biochemistry, University of Washington, Seattle, 98195, United States; Institute for Protein Design, University of Washington, Seattle, 98195, United States; Molecular Engineering Graduate Program, University of Washington, Seattle, 98195, United States

## Abstract

Programmable molecular biology increasingly requires strategies for converting engineered recognition or proximity modules into measurable outputs, particularly within transcriptional regulation, RNA imaging, and CRISPR-associated systems. Synthetic chemically induced dimerization (CID) systems provide a class of programmable recognition modules for such applications, yet generalized strategies for coupling structurally diverse CIDs to functional readouts remain limited. Here, we introduce a CID-to-output conversion strategy based on engineering of the linker-mediated coupling interface. Using single-fluorescent-protein sensors as an experimentally tractable optical model readout, we systematically varied paired N- and C-terminal linkers flanking circularly permuted green fluorescent protein (cpGFP) to map coupling landscapes across synthetic CID systems derived from combinatorial selection and computational protein design. The results revealed strong non-additive interactions across paired linkers and suggest that linker length is a first-order determinant of CID-to-output coupling. Across nanobody-, monobody-, and de novo-designed CID architectures, this framework yielded functional sensors with dynamic ranges up to 1270% and robust responses in mammalian cells. Together, this work demonstrates that effective CID-to-output conversion can be achieved by empirically mapping the linker-mediated coupling interface, providing a practical route for adapting synthetic CID to diverse programmable molecular readouts and nucleic-acid-associated synthetic biology systems

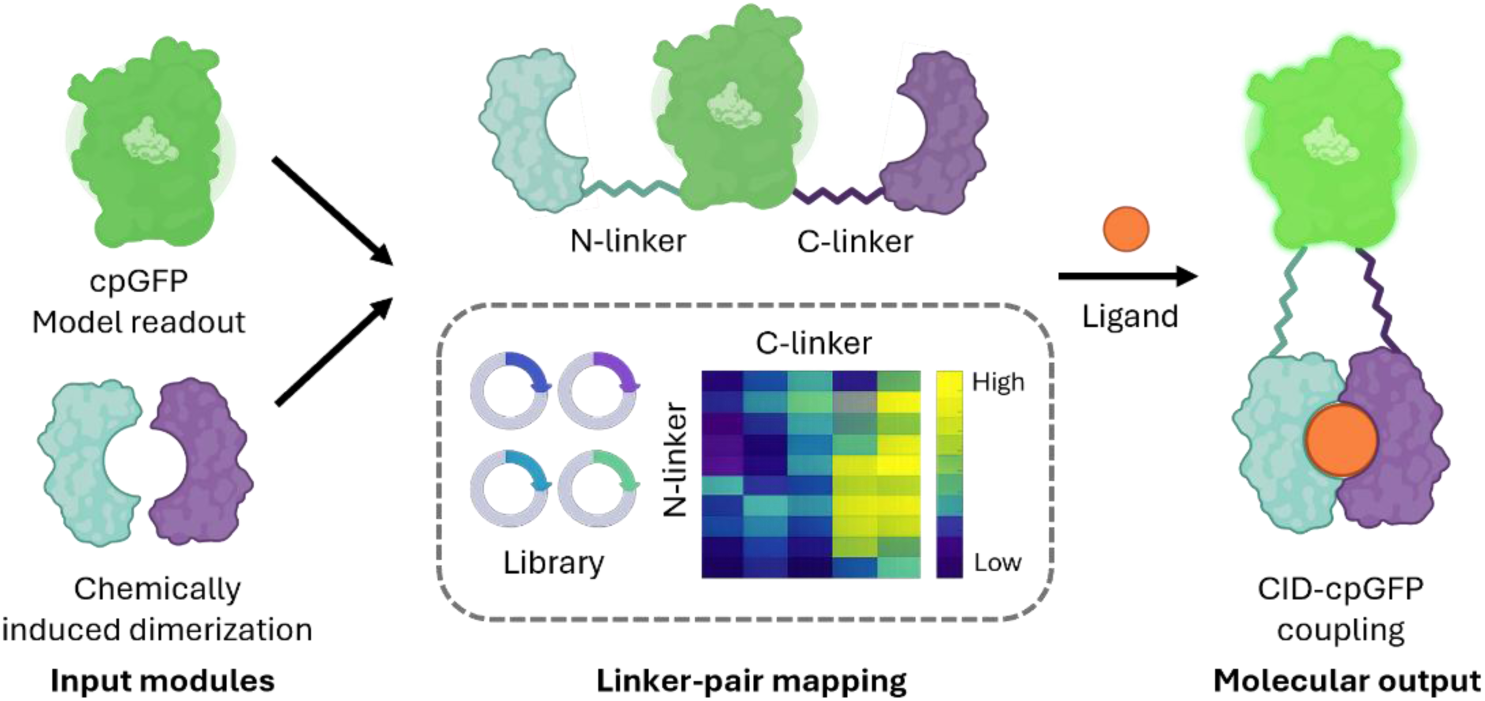

## Introduction

Engineered nucleic-acid-associated systems—including programmable transcriptional regulators, RNA-binding modules, and CRISPR-associated assemblies—often rely on modular protein components whose binding, proximity, or conformational states must be transduced into functional outputs [1–3]. However, converting these engineered molecular recognition events into robust downstream actions, such as transcriptional activation, molecular recruitment, or optical reporting, remains a challenge.

Synthetic chemically induced dimerization (CID) systems have emerged as powerful tools for programmable control and sensing in synthetic biology. By coupling ligand recognition to inducible molecular association, CIDs enable user-defined molecular sensing, regulation of localization, activity, signaling, and gene-regulatory programs [1, 4–11]. Recent advances in combinatorial selection [12–15] and computational protein design [16–19] have substantially accelerated the discovery of synthetic CID pairs responsive to diverse ligands, raising the prospect that such programmable molecular recognition modules can be generated on demand for a broad range of synthetic biological applications. However, CID discovery alone does not solve the downstream transduction problem: structurally diverse CIDs must still be coupled to output domains in geometries that permit both productive ligand-induced association and effective signal propagation [4, 5].

Genetically encoded fluorescent biosensors provide an attractive model system for studying this broader coupling problem. In particular, single-fluorescent-protein sensors based on circularly permuted fluorescent proteins (cpFPs), such as cpGFP offer straightforward optical readouts, high experimental accessibility, and direct sensitivity to local geometric perturbations [20–24]. Unlike split-reporter systems that primarily report proximity, cpFP-based sensors require productive mechanical coupling between ligand-induced structural rearrangements and the fluorescent chromophore environment, making them especially sensitive to linker architecture. They therefore provide a quantitative testbed for mapping how linker geometry governs recognition-to-readout transduction.

Previous studies demonstrated that linker optimization can strongly influence the performance of sensors based on naturally-occurring CID, as demonstrated by iFLinkC-enabled rapamycin-responsive FKBP12/FRB architectures [25, 26]. While these foundational studies highlighted the necessity of linker tuning, it remains unclear whether structurally diverse synthetic CIDs—particularly those discovered by combinatorial selection or de novo design—can be converted into functional readouts through a unified optimization strategy. Furthermore, it is unknown whether paired linker-length effects follow predictable, transferable rules or if they form highly system-specific, non-additive landscapes.

Here we introduce an engineering strategy for systematically coupling diverse synthetic CID systems to functional molecular outputs, using genetically encoded fluorescent biosensors as a model application. By treating N- and C-terminal linker lengths as coupled engineering variables and systematically exploring linker-length landscapes, we demonstrated that linker lengths can exert complex and strongly non-additive effects on coupling efficiency and established a “length-first” strategy for identifying permissive coupling geometries prior to sequence-level optimization. Using this framework, we engineered functional single-fluorescent-protein sensors from synthetic CID systems derived from nanobody-, monobody-, and de novo-designed architectures, validated their function in mammalian cells, and showed that structure-guided sequence refinement within a validated length regime can further improve performance. Together, these results establish linker-length landscape mapping as a practical engineering framework for converting structurally diverse synthetic CID systems into functional molecular readouts, providing a foundation for future adaptation of ligand-dependent interactions to broader sensing, regulation and nucleic-acid-associated synthetic biology applications.

## Material and Methods

### Reagents

Q5 Hot Start Master Mix (New England Biolabs, Ipswich, MA; cat. no. M0494); BbsI restriction enzyme (New England Biolabs; cat. no. R0539); T4 DNA Ligase (New England Biolabs; cat. no. M0202); EcoRI-HF (New England Biolabs; cat. no. R3101); NotI-HF (New England Biolabs; cat. no. R3189); AgeI-HF (New England Biolabs; cat. no. R3552); BglII (New England Biolabs; cat. no. R0144); B-PER Bacterial Protein Extraction Reagent (Thermo Fisher Scientific, Waltham, MA; cat. no. 78243); Axygen magnetic beads (Corning, Corning, NY; cat. no. MAG-PCR-CL-50); 0.45 µm syringe filter (Millipore/MilliporeSigma, Burlington, MA; cat. no. SLHV033RS); HisTrap HP 1 mL column (Cytiva, Marlborough, MA; cat. no. 17524701); HiTrap Desalting column (Cytiva; cat. no. 17140801); black 96-well microplate (Greiner Bio-One, Monroe, NC; cat. no. 655090); 8-well poly-D-lysine channel slides (Ibidi, Fitchburg, WI; cat. no. 80826); PEI Prime transfection reagent (Polyplus/Serochem; cat. no. prime-AQ100-100ML); high-glucose DMEM (Gibco/Thermo Fisher Scientific; cat. no. 11965092); penicillin-streptomycin (Gibco/Thermo Fisher Scientific; cat. no. 15140122); Live Cell Imaging Solution (Gibco/Thermo Fisher Scientific; cat. no. A14291DJ); Janelia Fluor 635 HaloTag Ligand (JF635; Promega, Madison, WI; cat. no. HT1050).

### Biological Resources

*E. coli* DH10B (standard laboratory strain). HEK293T cells (ATCC, Manassas, VA; cat. no. CRL-3216). pBAD expression vector (arabinose-inducible bacterial expression; addgene 54762). pCMV and pCMV-PRLSS vectors (mammalian cytoplasmic and ER lumen expression, respectively; from Sekhon et al. [27]). Modified pAEMXT vector (plasma membrane expression incorporating CD59 leader and anchor sequences; from Nasu et al. [28]). pAAV-synapsin-HaloCaMP1a-EGFP plasmid (cpHaloTag source; from Frei et al. [29]). CBD-responsive nanobody CID pair CA14/DB21 (from Kang et al. [13]). Computationally designed cortisol CID pair mhcy129/corD1 (from Chen et al. [16]). cpGFP crystal structure (PDB: 3WLC). CBD CID crystal structure (PDB: 7TE8).

### Statistical Analyses

High-throughput linker-library screening measurements were performed in technical triplicate and averaged prior to analysis. Unless otherwise noted, quantitative sensor characterization experiments, including dose–response measurements and EC₅₀ determinations, are presented as mean ± s.d. from three independent biological replicates. Biological replicates correspond to independently prepared protein samples or independent mammalian-cell transfections, as appropriate. Error bars represent standard deviation unless otherwise noted. Dose-response curves were fitted by nonlinear regression using a four-parameter logistic model: F = d + (a − d) / (1 + (x/c)^b), where a and d are the upper and lower asymptotes, c is the EC_50_, and b is the Hill coefficient; fitting was performed with MatplotLib. Non-additive interaction analysis used marginal-effect decomposition: the expected ΔF/F_0_ for each linker pair was calculated as the sum of the row-wise (N-linker) mean and column-wise (C-linker) mean, minus the grand mean across all combinations, with residuals defined as observed minus expected values. Pairwise structural RMSD comparisons used two-sided Mann-Whitney U tests (scipy.stats.mannwhitneyu; n = 5 Chai-1 predicted structures per linker variant; no correction for multiple testing applied). Significance thresholds: P < 0.05 (*), P < 0.01 (**).

### Novel Programs, Software, Algorithms

Non-additive interaction analysis and heatmap visualizations were implemented in Python. Code is available at https://github.com/davinan/gfp_sensor and deposited at Zenodo <u>10.5281/zenodo.18916934</u>. Plots were generated using Matplotlib (https://matplotlib.org). Statistical analyses used SciPy (https://scipy.org).

### Web Sites / Database References

AlphaFold3 (https://alphafoldserver.com; Abramson et al., Nature 630, 493–500, 2024). Chai-1 (https://chaistructure.com; Discovery et al., bioRxiv 2024.10.10.615955). RFDiffusion (https://github.com/RosettaCommons/RFdiffusion; Watson et al., Nature 620, 1089–1100, 2023). ProteinMPNN (https://github.com/dauparas/ProteinMPNN; Dauparas et al., Science 378, 49–56, 2022). RCSB Protein Data Bank (https://www.rcsb.org; Berman et al., Nucleic Acids Res. 28, 235–242, 2000). ImageJ (https://imagej.nih.gov; Schneider et al., Nat. Methods 9, 671–675, 2012). Matplotlib (https://matplotlib.org; Hunter, Comput. Sci. Eng. 9, 90–95, 2007). SciPy (https://scipy.org; Virtanen et al., Nat. Methods 17, 261–272, 2020).

### Linker length library and sensor library construction

N-terminal and C-terminal linkers with 253 bp overlap with cpGFP and a cpGFP fragment of 271 bp were ordered as eBlocks gene fragments (Integrated DNA Technologies, IDT). All starting DNA fragments used to generate the linker library are listed in Supplementary Table 1. To perform overlap-extension PCR, each combination of N-terminal and C-terminal linkers was mixed with cpGFP fragments in 96-well PCR plates. PCR reaction volume was 20 µL, including 10 µL Q5 Master Mix (New England Biolabs), and 1 µL of each DNA fragments (10 ng/µL each). PCR was conducted using a single-step protocol with 5 minutes annealing/extension at 72°C for 30 cycles. The PCR product was purified using magnetic beads (Axygen) and subsequently digested with BbsI (NEB) at 37°C for 30 min and deactivated at 75 °C for 20 min.

To generate the recipient vector, CID sequences were amplified by PCR and inserted into the pBAD vector via Gibson assembly, following the manufacturer’s instructions. A connector containing BbsI digestion sites and two glycine residues (DNA sequence: ggtggcttgtcttcatgttctctcccttggcagtaatagtgagaagacggcggt) was inserted between the CID pairs during the Gibson assembly. Overlaps with CID sequences are excluded from this connector, as they are CID-specific and must be designed individually for each CID pair.

To enhance intracellular stability of the CBD-responsive nanobody, framework mutations previously reported to improve cytosolic nanobody expression were introduced by ordering the mutated sequence as a gBlock gene fragment (Integrated DNA Technologies) and cloning it into the pBAD expression vector.

To generate the sensor library, linker length library fragments were ligated into the recipient vector in 96-well PCR plates using T4 DNA ligase (NEB) at a 3:1 insert-to-vector molar ratio, with incubation at RT for 15 min.

To validate library construction quality, colony PCR was performed on randomly selected clones using universal vector primers flanking the insert site (pBAD_seq_F: 5’-GCAACTCTCTACTGTTTCTC-3’; pBAD_seq_R: 5’-TTGACACTCATGGGTATGTATATC-3’). The expected amplicon size was approximately 1600 bp, with minor variation reflecting differences in linker length. Of 196 randomly selected clones, 190 contained inserts of the expected length, confirming high cloning fidelity. To verify sequence accuracy, 15 randomly selected variants and 9 variants with the longest linker combinations were submitted for Sanger sequencing (Azenta). Of 24 sequenced clones, 23 matched the intended amino acid sequences, demonstrating high concordance between the designed and cloned linker-length variants. Quality control was performed on the CBD sensor library and is representative of the general library construction workflow.

### Sensor library screening

The sensor library was individually transformed into DH10B bacteria using heat shock. Each clone was inoculated into 1 mL 2xYT media containing 100 μg/mL ampicillin in a 96-deep-well plate. Cultures were grown at 37βC until reaching an OD600 of ∼0.6, then induced with 0.02% arabinose and incubated overnight at 18βC. The cells were pelleted by centrifugation at 4,000 g for 5 minutes at room temperature and resuspended in 40 µL of B-PER bacterial protein extraction reagent (Thermo Fisher Scientific) mixed with 40 µL of PBS buffer. After incubating on a benchtop plate shaker at 1,500 rpm for 15 minutes at room temperature, 320 µL of PBS buffer was added to each well and thoroughly mixed. The mixture was centrifuged at 4,000 g for 5 minutes at room temperature, and the supernatant was carefully transferred to a 96-well plate, avoiding disturbance of the pellet. The fluorescence intensity and response of each clone were measured using a fluorescent plate reader (BioTek Synergy Neo2) at excitation/emission wavelengths of 488 nm/530 nm. 30 µM CBD, 1 mM MP and 1 µM cortisol were added to the lysate for each sensor library to measure sensor response. Technical triplicates were measured and averaged; plots were generated using MatplotLib.

### Protein expression & purification

All protein constructs were expressed in E. coli using a pBAD vector, with a C-terminal Avi-tag and His-tag and were subsequently purified by Ni-affinity chromatography. In brief, E. coli strain DH10B was transformed with the expression constructs using heat shock. Transformed cells were grown in 2xYT media containing 100 μg/mL ampicillin at 37βC until reaching an OD600 of ∼0.6. Protein expression was then induced with 0.02% arabinose, and cultures were incubated overnight at 18βC. Cell pellets from 1-liter cultures were harvested by centrifugation at 8000 g at 4 βC, resuspended in 15 mL ice-cold His-wash buffer (50 mM sodium phosphate, pH 8.0, 300 mM NaCl, 20 mM imidazole, 10% glycerol), and kept on ice. Bacterial cells were lysed using a Branson cell sonicator at 25% amplitude for 5 minutes with 5 seconds on and 5 seconds off, in an ice bath. The lysate was then centrifuged at 10,000 g, 4βC for 20 minutes to remove cell debris, followed by a second centrifugation of the supernatant at 20,000 g, 4βC for 30 minutes. The clarified supernatant was filtered through a 0.45 μm syringe filter (Millipore) at room temperature to remove any remaining particulates. The filtered supernatant was loaded onto a 1 mL HisTrap column (Cytiva) pre-equilibrated with His-wash buffer at a flow rate of 1 mL/min. The column was washed with 30 mL of His-wash buffer at 1 mL/min, and His-tagged proteins were eluted in a gradient manner, at ∼ 50% of elution buffer (50 mM sodium phosphate, pH 8.0, 300 mM NaCl, 250 mM imidazole, 10% glycerol). Eluted proteins were desalted using a 50 mL Hi-Trap desalting column (Cytiva), equilibrated with PBS buffer with 5% glycerol, at a flow rate of 5 mL/min. Desalted proteins were examined by SDS-PAGE.

### N12C2-cpHaloTag plasmid construction

The circularly permuted HaloTag (cpHaloTag) sequence was amplified from the pAAV-synapsin-HaloCaMP1a-EGFP plasmid [29] using primers flanking the cpHaloTag coding region. In parallel, the N12C2 sensor backbone was amplified using primers flanking the cpGFP core, thereby excluding the cpGFP optical domain. The cpHaloTag fragment was subsequently inserted into the amplified N12C2 backbone by Gibson assembly to generate the N12C2–cpHaloTag construct.

### Sensor *in vitro* characterization

To obtain excitation/emission spectra, 100 µL sensor protein solution of 1 µM was transferred to a black 96-well microplate (Greiner) and scanned by a fluorescent plate reader (BioTek Synergy Neo2). The wavelength range for cpGFP excitation/emission spectra is 400-520 nm/492-590 nm, with 2 nm increment. For cpHaloTag excitation/emission spectra, the range is 575-660 nm/640-720 nm, with 5 nm increment. To obtain spectra in the presence of the ligand, 30 µM CBD, 1 mM MP and 1 µM cortisol were added to protein solution of the corresponding sensor respectively.

To obtain dose-response curves, various concentrations of ligands were added to 100 µL sensor protein solution of 1 µM in a black 96-well microplate, mixed at 1000 rpm on a benchtop plate shaker for 30 seconds, and measured by a fluorescent plate reader. For CBD sensors, the CBD concentrations are 30, 3, 0.3, 0.03, 0.003, 3x10^-4^, 3x10^-5^ µM; for MP sensors, the MP concentrations are 1000, 333, 111, 37, 12.3, 4.1, 1.4 and 0 µM; for cortisol sensors, the cortisol concentrations are 10, 3.3, 1.1, 0.37, 0.12, 0.04, 0.01 and 0 µM.

To obtain basal fluorescence at various pH values, PBS buffers of pH 6, 6.4, 6.8, 7.2, 7.6, 8 were prepared. Sensor protein solution was diluted using these buffers to a final concentration of 1 µM, 100 µL of which was transferred to a black 96-well microplate and measured by a fluorescent plate reader. To obtain fluorescence response at various pH values, 30 µM CBD was added to the same wells and mixed at 1000 rpm on a benchtop plate shaker for 30 seconds before measurement. Readings before/after addition of CBD were used to calculate ΔF/F_0_, which is defined as F/F_0_-1. Both basal fluorescence and ΔF/F_0_ collected at different pH values were normalized against values at pH8. Data were analyzed using Excel. Briefly, the average and standard deviation for each triplicate were calculated; for dose response curves, nonlinear regression was performed using a four-parameter model (d + (a - d) / (1 + (x / c) ^b)), where a and d are the upper and lower asymptotes, c is the EC_50_, and b is the Hill coefficient. All plots were generated using MatplotLib. Unless otherwise stated, quantitative measurements are presented as mean ± s.d. from three independent biological replicates.

### Mammalian expression plasmids construction

To target cytoplasmic, ER lumen and cell surface expression, sensor variants were amplified by PCR and inserted into the pCMV vector [27], pCMV-PRLSS vector [27] and pAEMXT vector [28] using Gibson assembly. The pCMV vector was used for untagged cytoplasmic expression. pCMV-PRLSS vector adds the PRLSS leader sequence and KDEL ER-retention sequence to the protein. A modified pAEMXT vector incorporating a CD59 leader sequence and CD59 anchor sequence for plasma membrane display, as previously described [28], was used for surface expression. pCMV vector was digested with EcoRI and NotI (NEB); pCMV-PRLSS vector was digested with AgeI (NEB) and NotI; pAEMXT vector was digested with BglII (NEB) and EcoRI. PCR was conducted using a two-step protocol with 30-second annealing at 60°C and 2 minutes extension at 72°C for 30 cycles. All digestion reactions were carried out at 37βC for 30 minutes and deactivated at 75 βC for 20 minutes.

### Sensor validation in mammalian cells

HEK293T cells were cultured in high-glucose DMEM (Gibco) supplemented with 10% FBS and 1% penicillin-streptomycin (Gibco) at 37βC and 5% CO2. Cells were seeded at a density of 4x10^4^ in 8-well poly-D-lysine-coated channel slides (Ibidi) and transiently transfected with 0.3 µg of sensor plasmids using PEI Prime (Serochem) at a PEI ratio of 3:1 (µL: µg). After a 48-hour incubation period, transfected cells were washed with Live Cell Imaging Solution (Gibco) and transferred to a confocal microscope (Nikon) for imaging. Images were taken before the addition of various concentrations of ligand. For CBD sensor N12C2, to image the sensor response to ligands, Live Cell Imaging Solution supplemented with 30 µM CBD was added to each well in a remove-and-add manner, where the existing Live Cell Imaging Solution was carefully removed and replaced with Live Cell Imaging Solution containing the ligand. Cells were incubated for 3 minutes before images were captured. For the time-course experiments, 30 µM CBD and THC were added to the respective well using the same remove-and-add approach, and images were taken at 0, 30, 90, 150 and 300 seconds after the addition of the ligand.

Images were acquired using a Nikon Eclipse Ti inverted microscope equipped with a Yokogawa CSU-X1 spinning disk confocal unit and an Andor iXon Ultra 897 back-illuminated EMCCD camera. A 40× oil-immersion objective was used for all imaging. GFP and Cy5 filter sets were used for the 488-nm and 647-nm channels, respectively, with 488-nm and 640-nm laser lines for excitation. Images were acquired using NIS-Elements software (Nikon) and analyzed using ImageJ (NIH).

### Co-expression and imaging in mammalian cells

HEK293T cells were cultured in high-glucose DMEM (Gibco) supplemented with 10% FBS (Gibco) and 1% penicillin-streptomycin (Gibco) at 37βC and 5% CO2. Cells were seeded at a density of 4x10^4^ in 8-well poly-D-lysine-coated channel slides (Ibidi) and transiently co-transfected with 0.3 µg of each of the sensor plasmids using PEI Prime (Serochem) at a PEI ratio of 3:1 (µL: µg). After a 24-hour incubation period, transfected cells were labeled with 200 nM of Janelia Fluor® 635 HaloTag® Ligand (JF635, cat. no. HT1050, Promega). After a 48-hour incubation period, transfected cells were washed with Live Cell Imaging Solution (Gibco) and transferred to the microscope for imaging. Images were taken before the addition of ligand. To image the sensor response to ligands, Live Cell Imaging Solution supplemented with 30 µM CBD and 10 µM cortisol was added to each well using the same remove-and-add approach. Cells were incubated for 3 minutes before images were captured. Images were acquired and analyzed using the same microscope setup described above.

### Non-additive interaction analysis of linker combinations

To assess whether N- and C-terminal linker effects combine additively, we decomposed sensor performance into marginal and interaction-dependent components. For each linker combination, the observed ΔF/F₀ value was compared to an additive expectation constructed from the marginal effects of the corresponding N- and C-terminal linkers. Specifically, the expected ΔF/F₀ for a given linker pair was calculated as the sum of the row-wise (N-linker) mean and column-wise (C-linker) mean, minus the grand mean across all linker combinations. Residuals were then defined as the difference between the observed ΔF/F₀ and this additive expectation. Positive residuals indicate linker combinations that perform better than predicted from marginal effects alone, whereas negative residuals indicate underperformance. Residual values were visualized as heatmaps to highlight non-additive, interaction-dependent features of the linker–performance landscape.

### Structural prediction of cortisol sensor variants

AlphaFold3 (AF3) was used to model the de novo-designed cortisol chemically induced dimerization (CID) complex following the strategy of Chen et al [16]. Five models were generated, and the structure with the highest CID_pLDDT was selected as the CID reference for subsequent analyses.

Full-length sensor constructs consisting of mhcy129, circularly permuted GFP (cpGFP), and corD1 connected by glycine-serine (GS) linkers were predicted using Chai-1. Linker variants are denoted NxCx, where x indicates the GS linker length at the N- or C-terminal junction of cpGFP. For each construct (N2C2, N9C9, N9C13, N20C20), five structures were generated using Chai-1. Contact restraints derived from the cpGFP crystal structure (PDB 3WLC, chain A) were applied exclusively to the cpGFP region during modeling; no contact restraints were imposed on the CID. Cα-based Kabsch superposition was used to structurally align the Chai-1 predicted CID domain in the sensor construct to the AF3 predicted CID reference. Structural agreement was quantified as (i) CID Cα RMSD, computed over both CID chains and (ii) CID interface Cα RMSD, restricted to residues within 8 Å of an opposing chain in the AF3 predicted CID reference structure.

Pairwise comparisons of structural RMSD metrics across linker variants were performed using two-sided Mann-Whitney U tests. This nonparametric rank-based test was selected due to small sample size (n = 5 structures per variant). All pairwise contrasts were evaluated; P values are reported as P < 0.05 (*) and P < 0.01 (**). No correction for multiple testing was applied, given the exploratory nature of the structural comparisons. Statistical analyses were performed in Python using scipy.stats.mannwhitneyu.

### Linker computational design

Structures of the CID (PDB id: 7TE8) and of the cpGFP (PDB id: 3WLC) were used to find Cɑ atom contacts between residues. Contacts within 6 Å and 10 Å were used to provide restraints to Chai1, a general protein folding model, which predicted the structure of the full sequence (CA14_N-linker_cpGFP_C-linker_DB21), where CA14 and DB21 are the nanobody CBD-binding CID pair. A single random seed (63) was used to predict 5 structures, and the one with the highest overall pLDDT was kept as input to the design pipeline. The N-linker residue positions were then removed and RFDiffusion was used to re-generate backbone positions for these residues, resulting in 10 backbones. ProteinMPNN was used to generate 10 amino acid sequences for each generated backbone. The sequence design window was shifted to include the final residue of the CA14 nanobody (serine residue at position 119). This residue was included because it functions as a hinge governing CA14’s final orientation and may improve the stability of the ligand-bound sensor state. Finally, Chai1 was used to predict the structure of the generated sequences. Designs were filtered with low RMSD between the designed backbone and designed sequence predicted structure, and with high predicted CID pLDDT. Two designs were selected for in vitro characterization.

## Results

### Design of a systematic linker-length screening framework

Fluorescent readouts provide a scalable and quantitative format for measuring molecular signal transduction. We therefore selected CID-based fluorescent sensors as an experimental testbed for systematically dissecting how linker geometry influences recognition-to-readout coupling. Developing CID-based sensors requires coupling ligand-induced association of two independent binding partners to a cpGFP through peptide linkers, which imposes opposing constraints on dimerization and reporter coupling. Linker sequence can also affect reporter behavior, but jointly varying length and sequence creates a vast combinatorial search space [30]. We therefore asked whether linker length alone, in a sequence-controlled context, is sufficient to identify functional CID sensor architectures. To isolate length as the primary variable, we utilized glycine–serine (GS) linkers, which provide high conformational flexibility and minimal secondary structure propensity. We designed a sensor architecture with two GS linkers—designated the N-linker and C-linker—flanking the cpGFP core. By treating these as independent variables and varying each from 2 to 20 amino acids, we established a defined library of 361 unique linker-length combinations (Fig. 1C).

**Figure 1.**
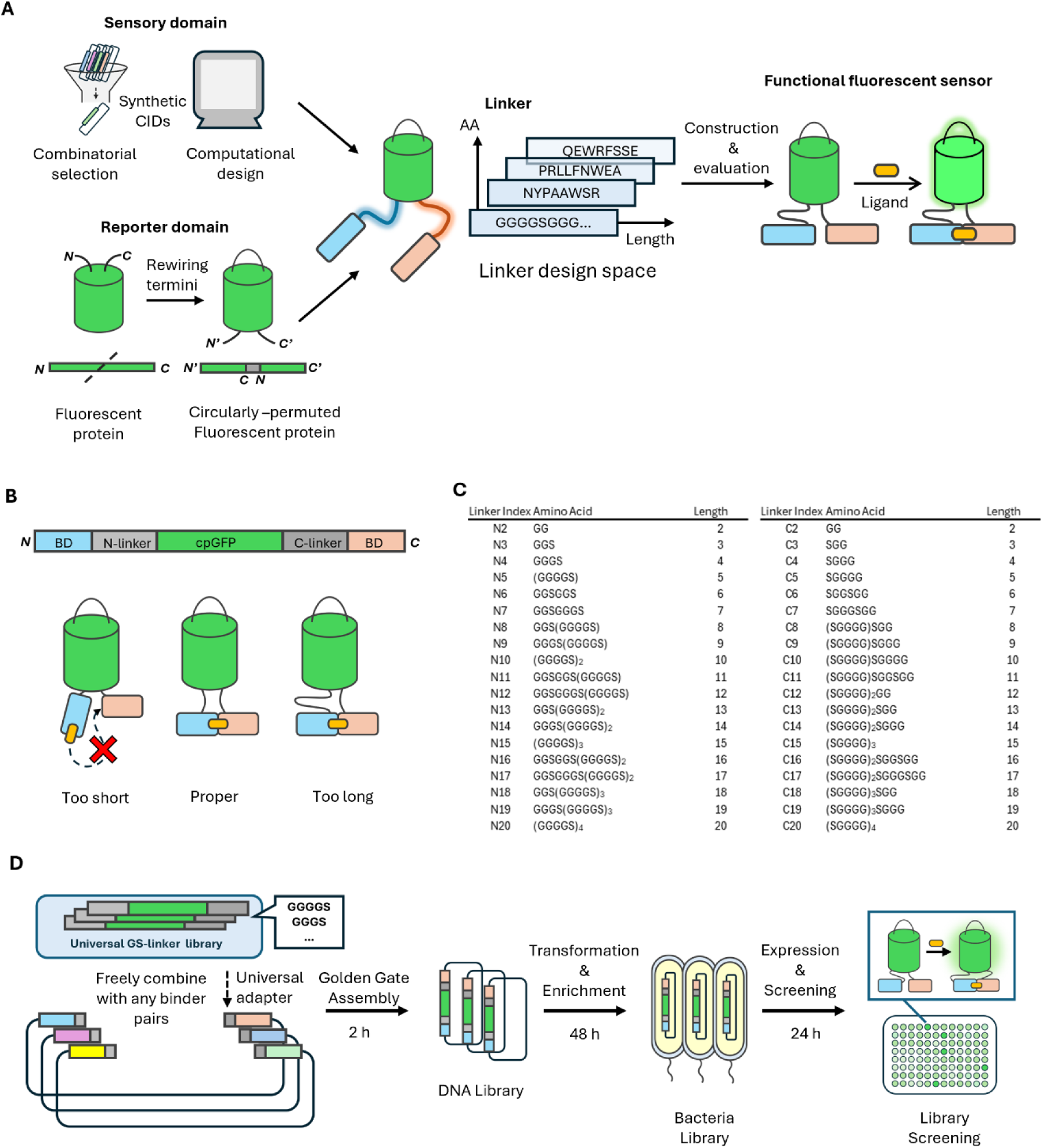
A modular framework for engineering CID-based fluorescent sensors. A, An overview of the framework. Synthetic chemically induced dimerization (CID) pairs derived from diverse sources, including combinatorial library screening and computational protein design, serve as interchangeable sensory domains. These CID pairs are mechanically coupled to circularly permuted optical reporter domains, such as cpGFP, through flexible peptide linkers of various lengths and sequences. The resulting CID-based sensors convert ligand-dependent dimerization into fluorescent readouts. B, Genetic architecture of CID-based fluorescent sensors with various linker lengths. Two flexible linkers flank the circularly permuted GFP (cpGFP) core and are referred to as the N-linker and C-linker; BD denotes the binding domain. Short linkers restrict domain mobility and can impede CID formation, whereas excessively long linkers reduce effective coupling between CID association and the fluorescent reporter. Intermediate linker lengths support both CID formation and ligand-dependent fluorescence modulation. C, Linker library composition. Both N-linker and C-linker consist of glycine–serine (GS) repeats with lengths ranging from 2 to 20 amino acids, yielding 361 unique linker-length combinations. D, Schematic of library construction and screening workflow. Insert fragments containing the cpGFP core flanked by variable linkers were generated by overlap extension PCR. Because the linker library encodes only the reporter and linkers, it is independent of CID identity and can be applied to different CID pairs. Insert fragments were ligated into CID-containing recipient vectors using Golden Gate cloning. The high insertion efficiency of Golden Gate assembly supports population-based bacterial screening without individual colony isolation.

To make the screening of the complete matrix practical, we implemented a modular cloning workflow (Fig. 1D). Variable linker fragments were generated by overlap-extension PCR and integrated into CID-containing recipient vectors using Type IIS Golden Gate assembly. Because assembly fidelity was consistently high, we optimized the workflow by expanding transformant populations under antibiotic selection rather than isolating individual colonies [31]. To validate the reliability of this population-based approach, we performed colony PCR on randomly selected variants and found 190/196 clones had insertion with expected length (Supplementary Fig. 1A). We subsequently performed Sanger sequencing of 15 randomly selected variants and 9 longest variants spanning the N-and C-linker ranges and observed high concordance with the intended designs, with 23 out 24 samples matching the expected amino acid sequences (Supplementary Fig. 1B). These quality-control results support the robustness of the population-based strategy. This matrix-based workflow enables the construction and initial screening of CID-sensor variants on a timescale of approximately four days, providing a rapid and portable route for translating new CID binders into genetically encoded sensor candidates.

### Systematic linker-length screening identifies a high-performance CBD sensor

To establish feasibility of the 2D linker-length framework and quantify how linker length shapes CID-based sensor performance, we first applied the library to a cannabidiol (CBD)-responsive CID pair [13]. This nanobody-based interaction pair was previously identified using COMBINES-CID, a combinatorial selection platform for high-affinity CID discovery. To ensure robust performance in the reducing environment of the cytosol, we first introduced framework mutations into the nanobody scaffold to enhance intracellular stability and expression [32] (Supplementary Fig. 2). We assembled the full 361-variant linker-length matrix with this CID pair and screened variants in bacteria, to quantify basal fluorescence and ligand-dependent responses; this lysate-based assay is intended primarily for relative ranking and hit identification. Basal fluorescence varied more than five-fold across the matrix, indicating that linker length alone can significantly perturb the reporter’s basal fluorescence independent of ligand binding (Fig. 2A). Notably, a subset of variants with a four–amino acid C-terminal linker (C4) consistently exhibited elevated basal fluorescence, regardless of N-linker length, suggesting that specific linker configurations can intrinsically favor a more fluorescent chromophore environment [33, 34](Fig. 2A, B). Analysis of the relationship between basal fluorescence and ligand-dependent response across the full matrix revealed that variants with elevated basal fluorescence tended to show reduced ligand-induced response amplitudes, suggesting that linker configurations that favor a more fluorescent apo state leave less headroom for ligand-induced fluorescence enhancement (Supplementary Fig. 3A).

**Figure 2.**
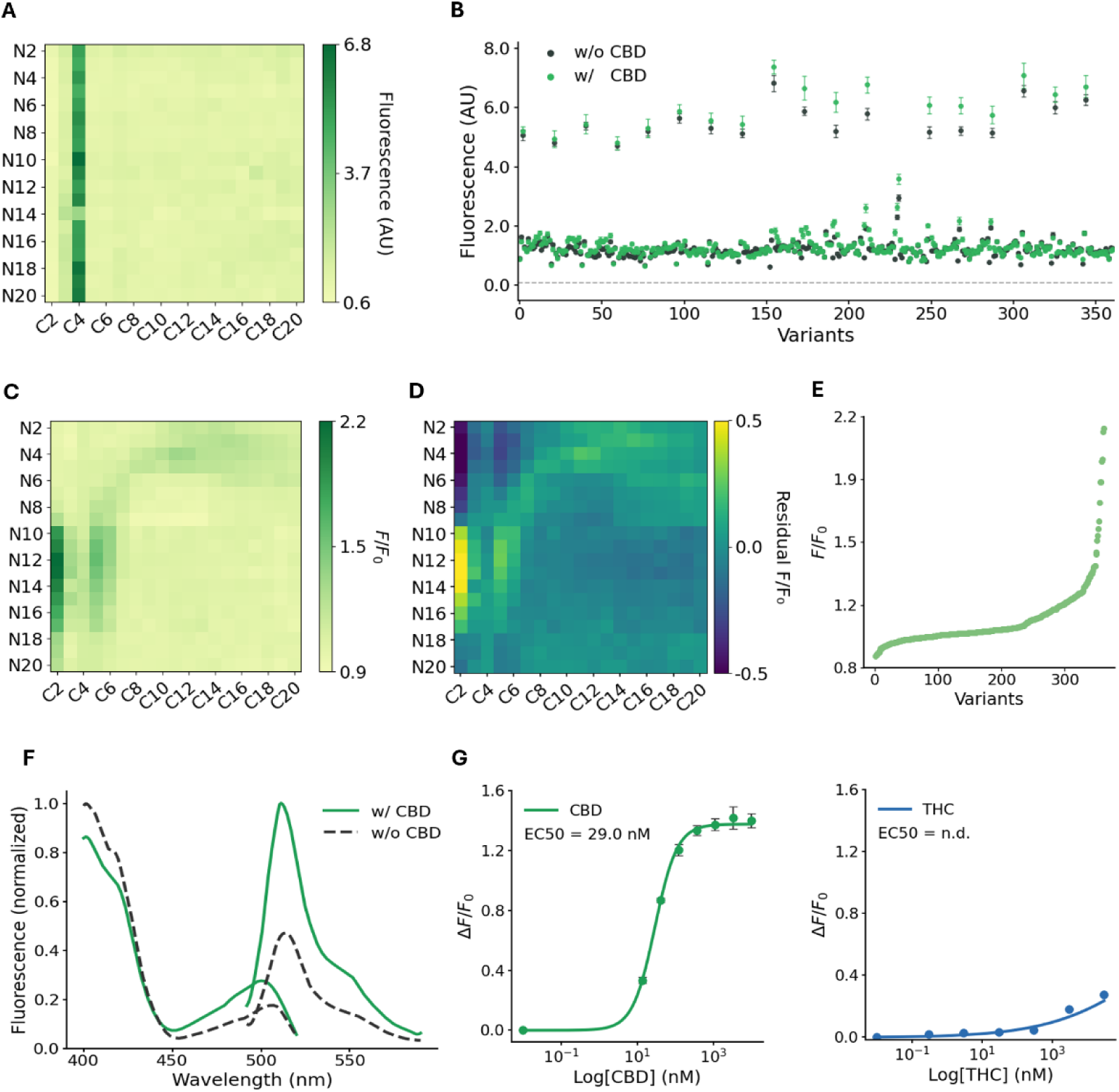
Linker library screening using a CBD-sensing CID pair. A, Heatmap of basal fluorescence (bacterial cell lysate) for all 361 linker variants of the CBD sensor. AU: arbitrary units. B, Fluorescence intensity of the same 361 variants before and after addition of saturating CBD ([CBD] = 30 µM). The dashed grey line indicates buffer background. Data are plotted as mean ± s.d. (n = 3 technical replicates). C, Heatmap of ligand-dependent fluorescence response of the same linker variants of the CBD sensor, expressed as F/F₀ (fluorescence at saturating ligand divided by basal fluorescence). D, Residual heatmap highlighting non-additive linker interactions. Residuals were calculated as observed F/F₀ minus an additive expectation defined as the sum of marginal N- and C-linker effects (row mean + column mean − grand mean). Positive values indicate linker combinations performing better than expected, while negative values indicate underperformance. E, Ranked response profile of all 361 CBD sensor variants, sorted by F/F₀. Screening data (A-E) were obtained from a single experiment with three technical replicates per variant. F, Excitation and emission spectra of the N12C2 sensor in the absence and presence of CBD ([CBD] = 30 µM). Primary λex = 402 nm, secondary λex = 502 nm and λem = 512 nm. Representative spectra measured from purified protein. G, Dose–response curves (purified protein) of the best-performing CBD sensor variant (N12C2) for CBD (left) and THC (right). ΔF/F₀ is defined as (fluorescence after ligand addition − basal fluorescence) / basal fluorescence. For CBD, EC_50_ = 29.0 nM and maximum ΔF/F₀ = 142%. Data represent mean ± s.d. from three independent biological replicates. EC_50_ is not determined for THC across the tested concentration range (up to 30 µM) in three independent biological replicates.

We next measured ligand-dependent responses upon addition of CBD. A broad distribution of signal modulation was observed. High-performance variants (ΔF/F₀ > 100%) clustered significantly around a 2-amino-acid C-terminal linker (C2) (Fig. 2C). In contrast, no high-performing variants were observed when both linkers exceeded 10 amino acids, consistent with reduced coupling at longer linker lengths.

To assess whether linker effects could be explained by independent contributions from the N- and C-linkers, we compared observed ΔF/F₀ values to an additive expectation derived from marginal effects of each linker length. This analysis revealed widespread non-additive behavior across the matrix, with both positive and negative deviations from additivity (Fig. 2D). Such cooperativity indicates that N- and C-linker lengths interact to shape sensor output, suggesting that the mechanical state of the reporter is a function of the combined linker geometry. Consequently, sensor performance cannot be reliably predicted or achieved through sequential one-dimensional optimization. Consistent with this, ranking all 361 variants by response magnitude revealed a broad and continuous distribution of performance rather than a small number of isolated optima (Fig. 2E), supporting systematic exploration of 2D linker-length landscape.

The best-performing variant, designated N12C2, was selected for further characterization (Supplementary Table 2). Notably, lead candidates were chosen solely on the basis of highest F/F₀, without explicit deprioritization of variants with high-baseline fluorescence. N12C2 exhibited a ligand-dependent increase in fluorescence, driven by a shift in the excitation equilibrium from 402 nm to 502 nm (Fig. 2F). This is consistent with a shift toward the deprotonated chromophore state, as previously observed in certain cpGFP - based sensors [35, 36]. We further examined pH dependence of basal fluorescence and CBD-induced signal change across a range of pH values (Supplementary Fig. 4A,B), and as expected for cpGFP-based sensors, N12C2 exhibited pH sensitivity [21, 22]. The sensor displayed a sigmoidal dose-response with an EC₅₀ = 29.0 nM and a maximum ΔF/F₀ = 142% (Fig. 2G). Crucially, N12C2 showed minimal response to the structurally related cannabinoid Δ9-tetrahydrocannabinol (THC), confirming that reporter fusion preserves the high specificity of the original nanobody binders (Fig. 2G). Together, these data provide proof-of-concept that 2D linker-length screening can identify CID-based single-fluorescent-protein sensors with robust ligand-dependent modulation from a defined length-only library.

### Cross-scaffold portability of linker-length landscape mapping

To evaluate whether the linker-length framework is portable across distinct synthetic CID scaffolds, we next applied it to a newly selected metorphamide-responsive CID pair. This MP-responsive pair was generated using the same combinatorial selection pipeline as the CBD system, but uses a monobody-based scaffold rather than a nanobody scaffold [37]. Because the focus of this study is CID-to-output coupling rather than de novo CID discovery, we used the MP pair here primarily as an orthogonal scaffold test case and do not extensively describe its selection or biochemical characterization. The CBD and MP systems also share a related geometric feature: in both cases, the ligand-binding pockets are oriented toward the N-termini of the binding partners. This provided an opportunity to ask whether the effective linker-length regime identified from the CBD landscape, particularly the enrichment of high-response variants in short C-linker configurations, could guide efficient screening of a distinct, newly selected CID architecture with comparable binding-pocket orientation. Guided by the CBD coupling landscape, in which high-responding designs were enriched in a short C-linker regime, we evaluated a targeted subset of 152 linker-length combinations (N2–N20 × C2–C9) for the newly selected monobody-based MP system. This subset was designed to test whether a linker-length regime identified from a nanobody-based CID could guide sensor construction for a distinct scaffold, rather than to exhaustively optimize or characterize the MP CID pair itself. Within this subset, basal fluorescence varied across linker combinations (Fig. 3A), and MP addition produced a broad distribution of ligand-dependent responses (Fig. 3B). As in the CBD system, analysis revealed pronounced non-additive dependence on paired N- and C-linker lengths (Fig. 3C), and ranking variants by response magnitude showed a broad performance distribution within the screened subset (Fig. 3D). As observed for the CBD sensor, variants with elevated basal fluorescence showed reduced ligand-induced response amplitudes (Supplementary Fig. 3B). From this subset screen, a lead variant N11C3 (Supplementary Table 2) emerged with robust fluorescence modulation and a maximum ΔF/F₀ of 1270% (Fig. 3E, F). Dose–response analysis showed a monotonic response below approximately 100 µM MP, followed by reduced signal at supersaturating ligand concentrations, consistent with a hook effect caused by competitive occupancy of the two binding partners and impaired productive dimerization [38]. Subsequent evaluation of the full linker-length matrix confirmed that superior variants were not enriched outside the targeted short-C-linker region (Supplementary Fig. 5), supporting the use of landscape-guided subset screening to recover high-performing designs in this scaffold context.

**Figure 3.**
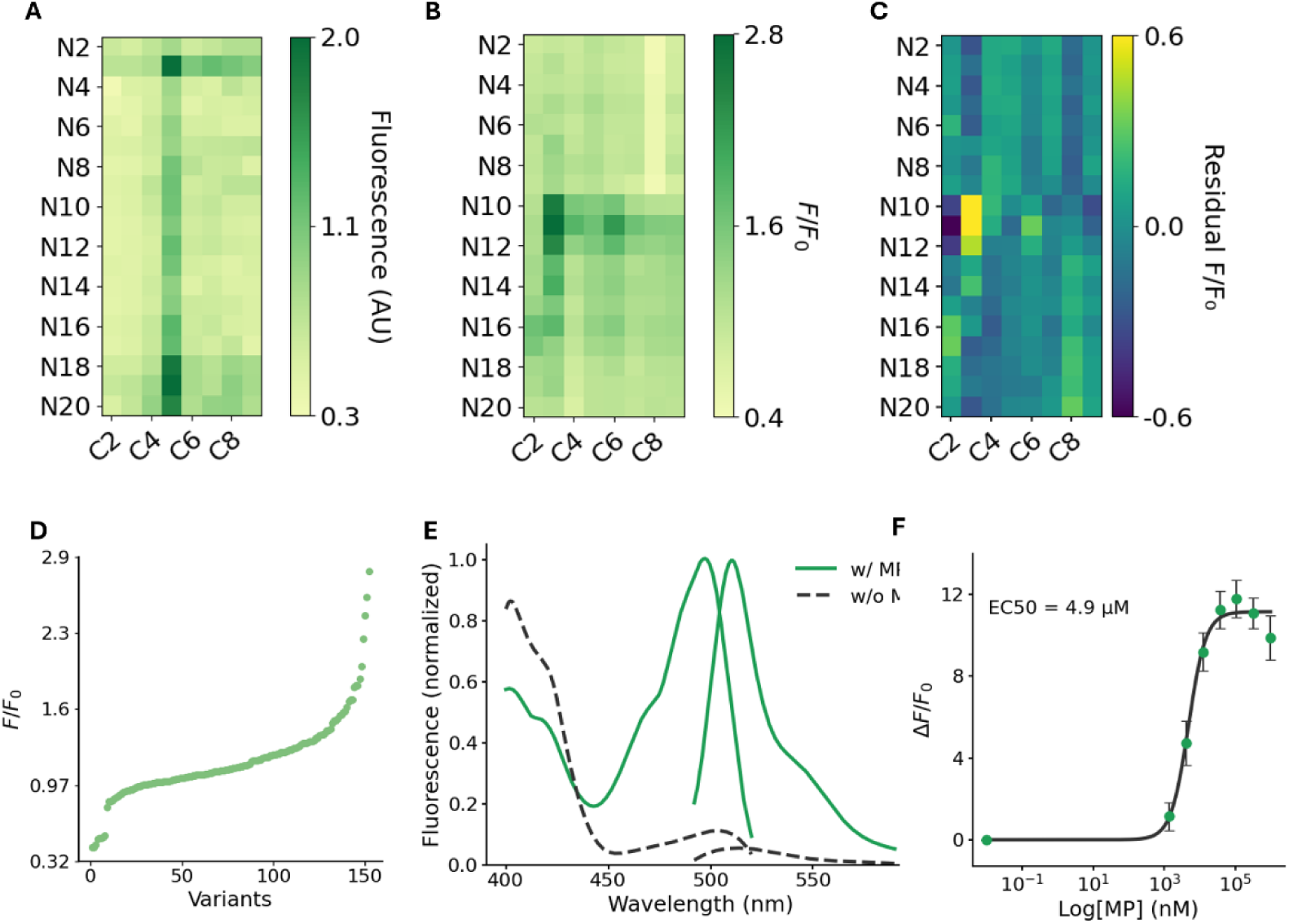
Linker library screening using a metorphamide-sensing CID pair. A, Heatmap of basal fluorescence (bacterial cell lysate) for 152 linker variants of the metorphamide (MP) sensor. N-linker: 2–20 aa; C-linker: 2–9 aa. AU: arbitrary units. B, Heatmap of ligand-dependent fluorescence response for the same 152 linker variants of the MP sensor, expressed as F/F₀ (fluorescence at saturating ligand divided by basal fluorescence). [MP] = 1 mM. C, Residual heatmap highlighting non-additive linker interactions. Residuals were calculated as observed F/F₀ minus an additive expectation defined as the sum of marginal N- and C-linker effects (row mean + column mean − grand mean). Positive values indicate linker combinations performing better than expected, while negative values indicate underperformance. D, Ranked response profile of all 152 MP sensor variants, sorted by F/F₀. Screening data (A-D) were obtained from a single experiment with three technical replicates per variant. E, Excitation and emission spectra of the N11C3 sensor in the absence and presence of MP ([MP] = 1 mM). Primary λex = 400 nm, secondary λex = 498 nm and λem = 510 nm. Representative spectra measured from purified protein. F, Dose–response curve (purified protein) of the best-performing MP sensor variant (N11C3). EC_50_=4.9 µM and maximum ΔF/F_0_ = 1270%. The dose–response curve shows a hook effect at high MP concentrations, consistent with competitive occupancy of the two binding partners. Data shown for [MP] = 0 to 1 mM. Data represent mean ± s.d. from three independent biological replicates.

### Structure-informed screening of a computationally designed cortisol CID

To evaluate whether the 2D linker-length framework extends beyond loop-based binders, we applied it to a cortisol-responsive CID pair generated by computational protein design [16]. Unlike the CBD and MP systems, which present binding interfaces through variable loops oriented toward the N-terminus, the cortisol CID has a distinct interface organization and complex geometry, providing an independent and more stringent evaluation of the framework’s generality.

Rather than screening the full 361-variant matrix, we used prior split-luciferase implementations of this CID as an empirical reference [16] to define a bounded search space spanning N2–N9 × C2–C13 (spanning 96 combinations). To assess geometric plausibility at the upper bound, we used AlphaFold3 [39] and Chai-1[40] to model the longest linker combination (N9C13); the resulting structure was consistent with accommodation of the CID interface without obvious steric obstruction (Supplementary Fig. 6). We then screened this structure-informed subset.

Within this restricted region, basal fluorescence varied across linker combinations (Fig. 4A), and addition of excess cortisol produced a broad range of ligand-dependent responses (Fig. 4B). As in the nanobody- and monobody-based systems, analysis revealed pronounced non-additive interactions between N- and C-linker lengths (Fig. 4C), and ranking variants by response magnitude showed a broad performance distribution (Fig. 4D). Similarly, variants with elevated basal fluorescence showed compressed ligand-dependent responses (Supplementary Fig. 3C). The best-performing design, N5C8, exhibited ligand-dependent spectral modulation (Fig. 4E) and a sigmoidal dose response with EC₅₀ = 20.3 nM and a maximum ΔF/F₀ = 364% (Fig. 4F), with a hook effect apparent at supersaturating cortisol concentrations [16]. The sensor response is monotonic below ∼1 µM cortisol. Together, these results show that bounded, structure-informed exploration of 2D linker-length space can identify a functional cortisol sensor architecture and that coupled linker dependence persists in a computationally designed CID system.

**Figure 4.**
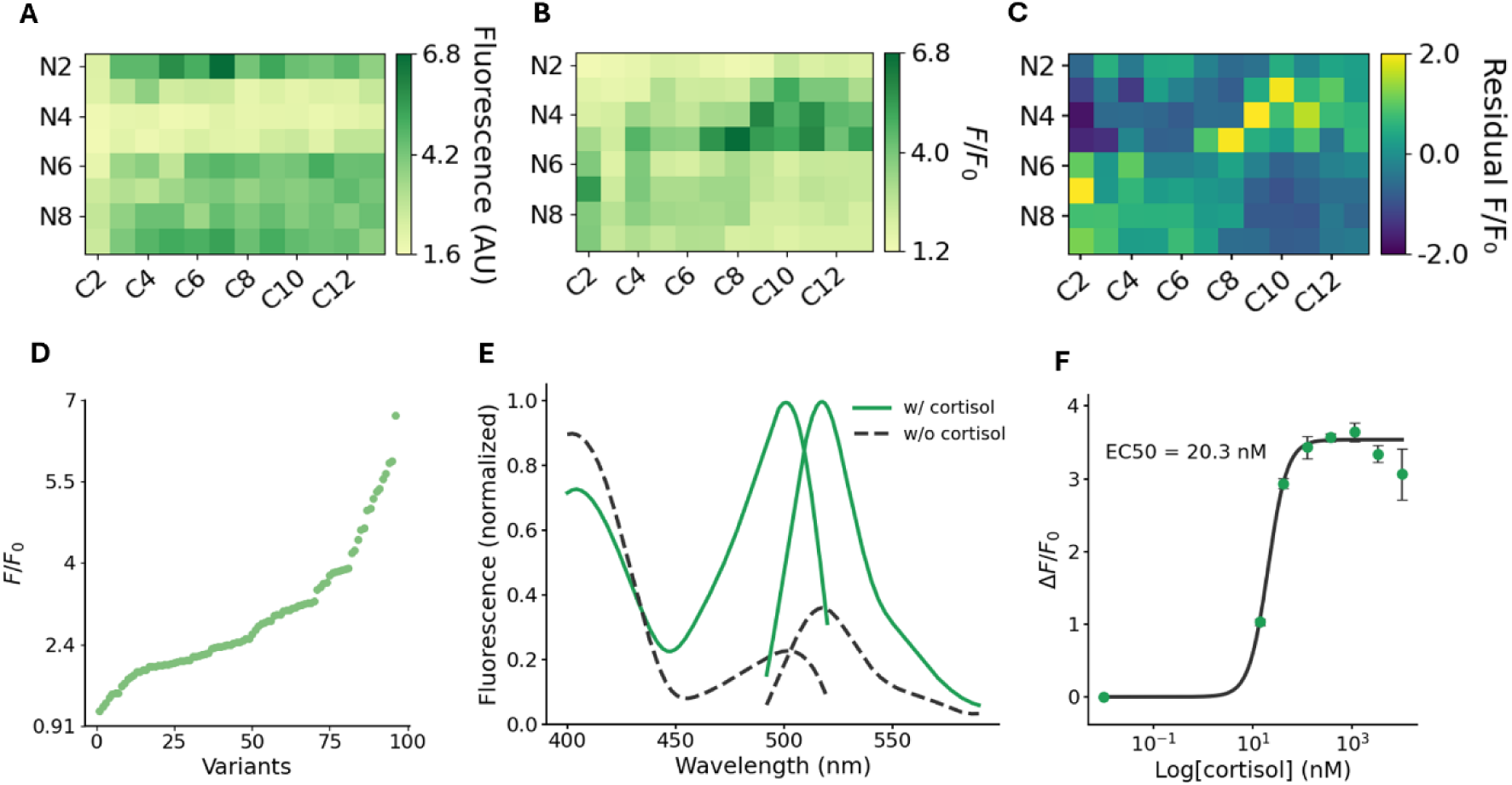
Linker library screening using a cortisol-sensing CID pair. A, Heatmap of basal fluorescence (bacterial cell lysate) for 96 linker variants of the cortisol sensor. N-linker: 2–9 aa; C-linker: 2–13 aa. AU: arbitrary units. B, Heatmap of ligand-dependent fluorescence responses for the same 96 linker variants, expressed as F/F₀ (fluorescence at saturating ligand divided by basal fluorescence). [cortisol] = 1 µM. C, Residual heatmap highlighting non-additive linker interactions. Residuals were calculated as observed F/F₀ minus an additive expectation defined as the sum of marginal N- and C-linker effects (row mean + column mean − grand mean). Positive values indicate linker combinations performing better than expected, while negative values indicate underperformance. D, Ranked response profile of all 96 cortisol sensor variants, sorted by F/F₀. Screening data (A-D) were obtained from a single experiment with three technical replicates per variant. E, Excitation and emission spectra of the N5C8 sensor in the absence and presence of cortisol ([cortisol] = 1 µM). Primary λex = 402 nm, secondary λex = 502 nm and λem = 518 nm. Representative spectra measured from purified protein. F, Dose–response curve (purified protein) of the best-performing cortisol sensor variant (N5C8). EC_50_ = 20.3 nM; maximum ΔF/F_0_ = 364%. The dose–response curve shows a hook effect at supersaturating cortisol concentrations. Data shown for [cortisol] = 0 to 10 µM. Data represent mean ± s.d. from three independent biological replicates.

### Mammalian-cell validation across diverse synthetic CID architectures

We next asked whether sensor variants identified by 2D linker-length screening remain functional in mammalian cells and retain modularity for multichannel imaging. We first characterized the lead CBD sensor, N12C2, which was expressed in HEK293T cells and targeted to the plasma membrane (PM), cytoplasm, or endoplasmic reticulum (ER) lumen. In each subcellular context, addition of CBD produced a ligand-dependent fluorescence increase (Fig. 5A, Supplementary Fig. 7A), indicating that linker-length–derived architectures identified in bacterial screening can function upon mammalian expression and subcellular targeting.

**Figure 5.**
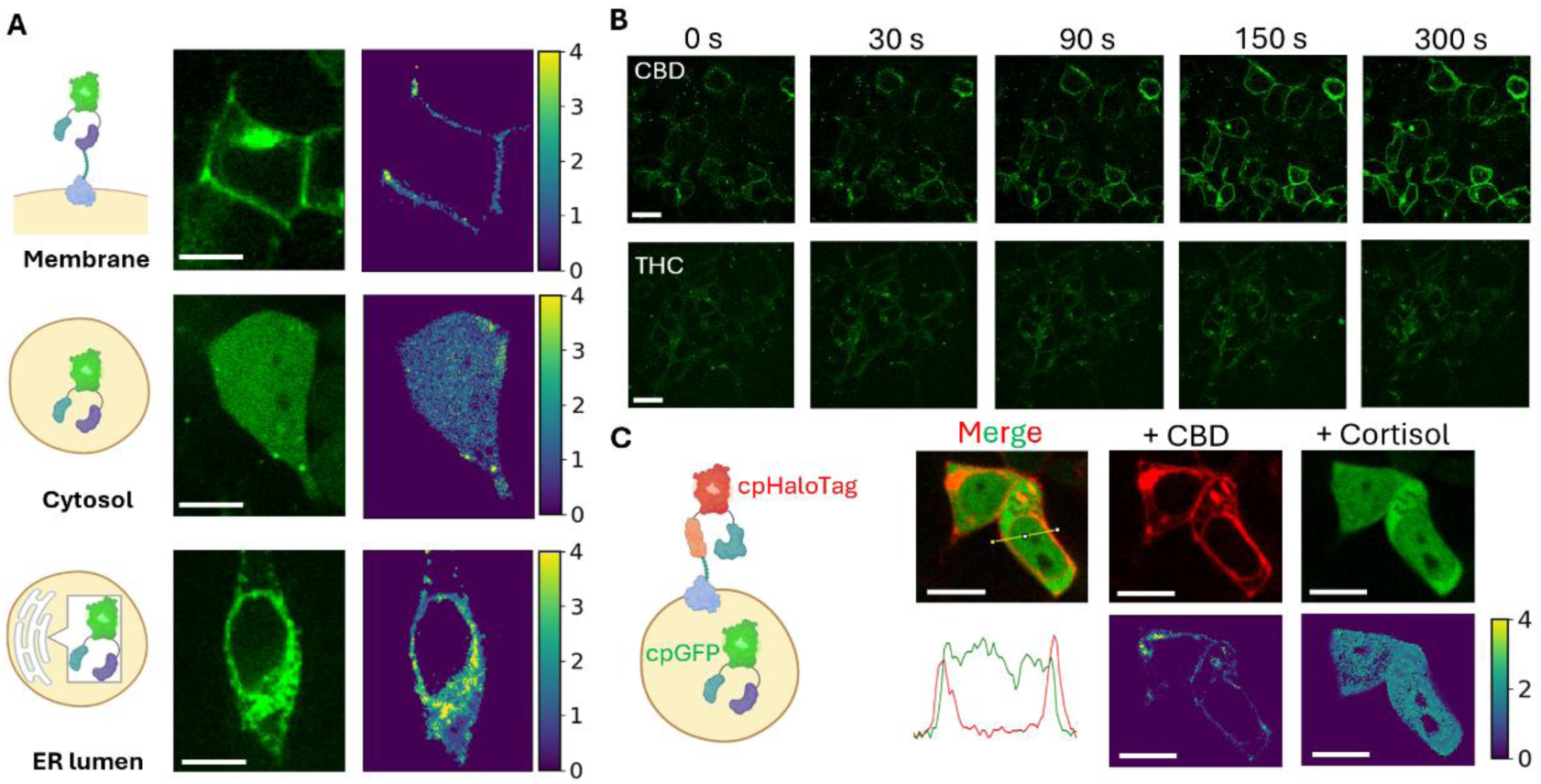
Cellular expression, ligand responsiveness, and multiplexed imaging of CID-based biosensors. A, Representative fluorescence images showing N12C2 expressed on the plasma membrane (top), in the cytoplasm (middle), and in the endoplasmic reticulum lumen (bottom) of HEK293T cells before ligand addition (left), after addition of 30 µM CBD (middle), and ΔF/F₀ ratio image (right). Color scale bar, ΔF/F₀. Scale bar, 20 µm. B, Representative time-lapse fluorescence images of surface-displayed N12C2 following addition of 30 µM CBD (top), 30 µM Δ⁹-tetrahydrocannabinol (THC, bottom) over 300 s. Scale bar, 40 µm. C, Representative fluorescence images showing the overlay (left), N12C2-cpHaloTag CBD sensor (middle, 647-nm channel), and N5C8 cortisol sensor (right, 488-nm channel) in HEK293T cells. The line-scan profile shows fluorescence intensity across both channels along the yellow line, confirming co-expression of both sensors within the same cells. For each channel, images show fluorescence after addition of 30 µM CBD and 10 µM cortisol (top), and ΔF/F₀ ratio image (bottom). Color scale bar, ΔF/F₀. Scale bar, 20 µm. Fluorescence images were acquired using identical settings within each channel. All experiments were repeated independently three times with similar results.

We then evaluated cellular specificity, kinetics, and dose dependence using the surface-displayed sensor. The sensor demonstrated high specificity, showing no measurable response to 30 µM Δ9-tetrahydrocannabinol (THC), a structurally related cannabinoid, whereas maintaining a strong response to CBD (Fig. 5B). Under the tested delivery conditions, CBD responses approached steady state on the order of minutes (Supplementary Fig. 7B). Titration of PM-targeted N12C2 across increasing CBD concentrations revealed a concentration-dependent response with an apparent EC₅₀ = 197 nM and a maximum ΔF/F₀ = 103% (Supplementary Fig. 7C). The apparent cellular EC₅₀ was higher than that measured with purified protein, which may be due to partitioning of CBD into the lipid bilayer or constraints on dimerization imposed by membrane tethering.

To determine whether mammalian-cell activity extends beyond the nanobody-based CBD sensor, we next evaluated lead sensors derived from the monobody-based MP CID and the de novo-designed cortisol CID. The MP sensor (N11C3), targeted to PM retained robust ligand-dependent fluorescence modulation in HEK293T cells, with a maximum ΔF/F₀ of 420% and an apparent EC₅₀ of 2.4 µM (Supplementary Fig. 8A, B). The cortisol sensor N5C8, expressed cytoplasmically, also showed clear ligand-dependent activity, with a maximum ΔF/F₀ of 140% and an apparent EC₅₀ of 145.2 nM (Supplementary Fig. 8C, D). Together with the CBD sensor, these results show that linker-length-derived readouts from nanobody-, monobody- and de novo-designed CID architectures can remain functional in mammalian cells, supporting the mammalian-cell compatibility of the coupling framework across distinct synthetic CID systems. For both the MP and cortisol sensors, measurements in cells showed reduced dynamic range and increased apparent EC₅₀ relative to purified protein measurements, indicating that cellular context, subcellular localization and ligand availability can influence quantitative sensor performance.

Finally, we tested whether 2D linker-length screening–derived sensor variants support multiplexed imaging through optical-domain swapping. We co-expressed the cortisol sensor N5C8 with a modified CBD N12C2 sensor in which the cpGFP domain was replaced by cpHaloTag labeled with JF635 [29, 41]. This configuration enabled spectrally orthogonal readouts in the 488-nm and 647-nm channels, respectively. The resulting cpHaloTag construct retained ligand-dependent modulation, suggesting compatibility of the 2D linker-length screening framework with alternative fluorescent reporters. Representative images showed simultaneous, ligand-dependent responses in both channels following addition of 30 μM CBD and 10 μM cortisol (Fig. 5C), demonstrating that CID-based sensors with distinct optical reporters can be co-expressed and used for multichannel imaging in single cells. Together with the cellular characterization of CBD, MP and cortisol sensors, these results support the mammalian-cell compatibility of linker-length-derived CID readouts and provide an initial demonstration that selected CID–linker architectures can be adapted to an alternative reporter format.

### Sequence level refinement within a validated length regime

Having established 2D linker-length screening as a robust and generalizable method to identify functional CID-based sensors, we next asked whether sequence-level refinement within an empirically validated length configuration could further enhance performance. Rather than revisiting linker length, we treated the optimized length regime as a fixed parameter and tested whether linker sequence changes could further enhance mechanical coupling between the CID pair and the reporter.

Using the lead CBD sensor variant N12C2 as proof of concept, we first used Chai-1 to predict the structure of the assembled N12C2 sensor from the cpGFP crystal structure and the CID complex, providing a structural basis for linker redesign (Supplementary Fig. 9). We then redesigned the N-terminal linker using RFdiffusion [42]/ProteinMPNN [43]-guided modeling to bias amino acid compositions while strictly preserving the validated linker lengths (Fig. 6A, B). This redesign was intended to alter linker backbone geometry and sequence composition within the validated length regime to improve CID-to-reporter coupling. Because the design procedure does not directly model cpGFP chromophore protonation, maturation or fluorescence output, we interpreted these variants as empirical sequence refinements rather than mechanistic optimization of chromophore photophysics.

**Figure 6.**
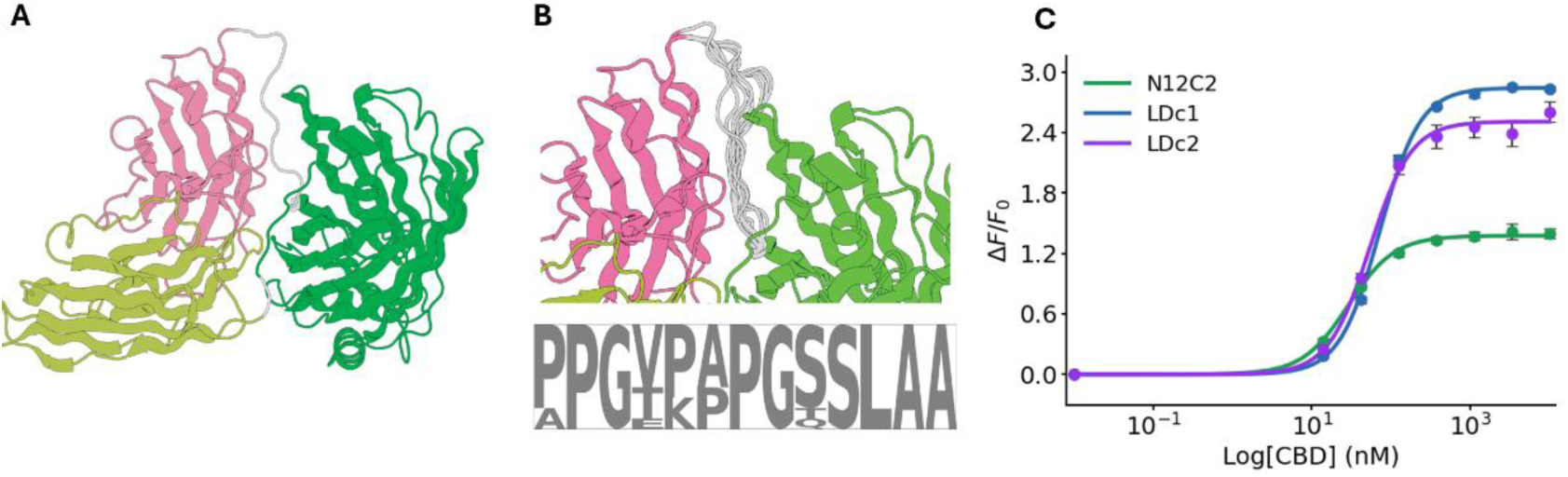
Structure-guided computational redesign of sensor linkers. A, Representative Chai1-predicted structural model of the CBD sensor (N12C2), illustrating the spatial arrangement of the binding domains, linkers, and cpGFP core. The model is used as a structural basis for linker redesign. B, Structural diversity of RFdiffusion-generated linker backbones of fixed length (top), and corresponding sequence logo summarizing position-wise amino-acid preferences of the N-linker post ProteinMPNN design (bottom). C, Dose–response curve (purified protein) of the best-performing CBD sensor N12C2 and its two designed linker variants, LDc1 and LDc2, selected as the top-ranked designs from RFdiffusion/ProteinMPNN. ΔF/F₀ is defined as (fluorescence after ligand addition − basal fluorescence) / basal fluorescence. The N12C2 dose–response curve is reproduced from Fig. 2G for comparison; LDc1: EC_50_ = 72.2 nM; maximum ΔF/F_0_ = 283%; LDc2: EC_50_ = 53.9 nM; ΔF/F_0_ = 260%. Data represent mean ± s.d. from three independent biological replicates.

The two top-ranked redesigned variants, LDc1 and LDc2 (Supplementary Fig. 10 ), showed increased maximal response, reaching ΔF/F₀ = 260–283% compared to the parent GS-linker design (ΔF/F₀ = 129%) (Fig. 6C). The EC₅₀ values remained similar, indicating that sequence-level changes primarily affected dimerization-to-fluorescence coupling efficiency rather than ligand recognition. Although we cannot exclude contributions from altered folding, maturation or basal reporter state, these results show that linker sequence can further modulate sensor performance within an empirically validated length configuration. Together, these data support a hierarchical linker-engineering workflow in which two-dimensional linker-length mapping first identifies permissive coupling geometries, followed by targeted sequence-level refinement to improve output performance within the selected length regime.

## Discussion

The rapid expansion of synthetic chemically induced dimerization (CID) technologies through combinatorial selection and computational protein design is creating new opportunities for ligand-controlled molecular engineering. However, as the number and diversity of available CID pairs increases, an important bottleneck is shifting from the discovery of ligand-dependent recognition modules to the challenge of coupling those modules to functional molecular outputs. CID systems can, in principle, be linked to many downstream mechanisms, including optical readouts, transcriptional regulators, RNA-associated systems, CRISPR-based assemblies and other programmable synthetic biology architectures. However, the physical interface that converts ligand-induced dimerization into an output response often remains difficult to design rationally. In the present study, we used single-fluorescent-protein biosensors as an experimentally tractable model system to investigate general principles governing CID-to-output coupling.

CID-based sensor engineering presents a distinct design problem from classical allostery-based cpFP sensor development. Because CID coupling depends on ligand-induced dimerization rather than conformational change within a single allosteric protein, the established heuristics that guide cpFP insertion in allostery-based sensors are not transferable. Previous studies demonstrated that linker optimization can substantially influence CID-based single-fluorescent-protein sensor performance, including rapamycin-responsive FKBP12/FRB architectures engineered using iFLinkC [25, 26]. These studies established the feasibility of CID-based single-FP sensing and highlighted the importance of linker engineering, but did not systematically investigate how coupled linker lengths shape multidimensional performance landscapes across diverse CID systems. Here we show that systematic, two-dimensional exploration of 2D linker-length space provides a practical strategy to optimize linker lengths for CID-based sensors. Across three CID systems spanning combinatorial selection and computational design, this workflow yielded single-fluorescent-protein sensors with large ligand-dependent fluorescence changes up to ∼1270% ΔF/F₀, comparable to many reported cpFP-based sensors [21].

A central observation across all tested CIDs is the non-additive dependence of sensor performance on paired N- and C-linker lengths. Rather than operating independently, the two linkers act in concert to transduce CID dimerization into fluorescence modulation of the reporter, such that the optimal choice of one linker depends on its paired partner. This behavior indicates that optimizing linker lengths jointly, rather than independently, is necessary to accurately map the geometric coupling in composite fusions spanning two associating proteins. These CID-specific coupling landscapes clarify what is portable in the present framework. The reusable component is not a universal linker sequence or length, but the screening strategy itself: a modular linker library and matrix-based cloning workflow that can be applied to each new CID pair. Because optimal linker configurations remain architecture-specific, each CID system requires empirical mapping of its own coupling landscape. While full-matrix 2D linker-length screening provides the most complete picture of coupling behavior, our results also suggest practical ways to reduce experimental burden when prior structural or empirical knowledge is available. For the monobody-based CID, subset screening guided by the CBD-sensor coupling landscape recovered a high-performance variant, and full-matrix evaluation shows no enrichment of superior variants outside the subset region. For the computationally designed cortisol CID, prior implementation constraints combined with structural modeling defined a bounded search space that still yielded a high-performing sensor. Together, these findings suggest flexible approaches to reducing experimental burden: a full-matrix 2D linker-length screen for one CID pair can inform subset screening for structurally related CID pairs, while structural knowledge can narrow the search space for architecturally distinct ones.

The mammalian-cell data support the practical utility of linker-length-derived CID readouts. Lead sensors from nanobody-based CBD, monobody-based MP and de novo-designed cortisol CID architectures all retained ligand-dependent activity in HEK293T cells. The CBD sensor functioned across multiple subcellular localizations and preserved discrimination against THC, while the MP and cortisol sensors also showed robust cellular responses. However, apparent EC₅₀ values and dynamic ranges differed from purified-protein measurements, emphasizing that quantitative sensor behavior must be validated in the intended cellular context. As an initial proof of concept for reporter adaptability, replacement of cpGFP with cpHaloTag in the CBD sensor showed that at least one optimized CID–linker architecture can tolerate an alternative optical reporter format.

Our results also support a hierarchical linker-engineering workflow in which linker length is optimized before linker sequence. Initial GS-linker screens isolated length as the primary variable and avoided the much larger search space created by simultaneous variation of length and sequence. After identifying the N12C2 length configuration for the CBD sensor, RFdiffusion/ProteinMPNN-guided sequence refinement within this fixed length regime increased response amplitude without major EC₅₀ shifts. These data suggest that linker sequence can further tune output efficiency after a productive length regime has been identified, although the redesign should be viewed as an empirical refinement rather than a mechanistic optimization of cpGFP photophysics.

Several limitations of the present study merit consideration. First, while bacterial screening provided a practical and efficient route to identify functional linker-length configurations, both the apparent affinity and dynamic range of the CBD sensor were reduced in mammalian cells relative to purified protein measurements, suggesting that in vitro characterization may not fully predict quantitative performance in cellular contexts. Factors such as macromolecular crowding, subcellular compartmentalization, membrane partitioning, and proteolytic degradation may influence sensor behavior in ways that are not captured in bacterial screens, and cellular validation therefore remains an essential step in the workflow. For sensors exhibiting a hook effect, such as the MP and cortisol sensors, the monotonic working range should be established empirically before use. We also note that lysate-based dynamic range measurements did not quantitatively predict purified-protein performance, consistent with the variable expression and complex background of crude lysate; for peptide-responsive systems such as the MP sensor, ligand instability or degradation in lysate may further compress the observable response. Lysate screening is therefore best interpreted as a tool for rank-ordering and hit identification rather than quantitative benchmarking. Second, the current framework was validated across three CID systems of relatively modest size; whether the same linker-length dependencies hold for larger or more structurally complex CID components — such as those involving membrane proteins or multidomain scaffolds — remains to be established. Third, this study focused exclusively on GS linkers as a sequence-controlled, flexible baseline; other linker compositions with distinct rigidity or secondary structure propensity were not systematically explored. Although sequence-level refinement using RFdiffusion/ProteinMPNN partially addresses this, a broader exploration of linker composition may reveal additional performance gains in future work. Finally, the present study used fluorescent biosensors as the main model output. Although this is a useful and stringent readout for measuring coupling efficiency, direct extension to non-optical outputs, such as transcriptional activation, RNA localization or CRISPR-associated recruitment, will require output-specific validation.

Overall, this work establishes linker-length landscape mapping as a practical strategy for converting synthetic ligand-dependent interactions into functional molecular readouts. The key principle is not that one linker design is universally transferable, but that systematic mapping can identify permissive coupling geometries for each CID architecture. This length-first, sequence-second workflow provides a scalable starting point for adapting newly discovered synthetic CID systems to programmable sensing, regulation and nucleic-acid-associated synthetic biology applications.

## Data Availability

The raw fluorescence screening data, Python analysis scripts and codes are available at Zenodo <u>10.5281/zenodo.18916934</u>. Structural coordinates used in this study are available from the RCSB Protein Data Bank under accession codes 3WLC and 7TE8.

## Supplementary Data

Supplementary Data are available at NAR Online.

## Acknowledgements

*Author contributions*: L.G. conceived the original project, supervised the study, provided resources, and acquired funding. S.K., Y.P. and L.G. conceptualized the study. Y.P. and S.K. performed the experiment and developed the methodology. D.N.A. and Y.P. performed linker computational design, and analyzed the data. Y.P performed the cell experiments, and analyzed the data. The manuscript draft was written by Y.P. and S.K.. Y.P., S.K., D.N.A., Y.Y., F.D. and L.G. reviewed and edited the manuscript. All authors have given approval to the final version of the manuscript. We thank Dr.Robert E. Campbell at the University of Tokyo for in-depth discussions.

The authors used ChatGPT (model 5.5) to improve the manuscript’s language and readability. The final content was reviewed and approved by the authors, who take full responsibility for the text.

The graphic abstract and schematics in Figure 5 were created in BioRender. Sun, L. (2026) https://BioRender.com/llglisc

## Funding

This work was supported by grants from the U.S. National Institutes of Health (NIH) (1R35GM128918, R21DA051555, R21DA051194, and R61/R33DA051489) to L.G.

## Conflict of interest disclosure

A patent related to this work has been filed by the University of Washington.

## Notes

### Competing Interest Statement

The authors have declared no competing interest.

