## Supplementary Material for "Linker-Length Landscape Mapping Enables Coupling of Diverse Synthetic Chemically Induced Dimerization Systems to Molecular Readouts"

**A**

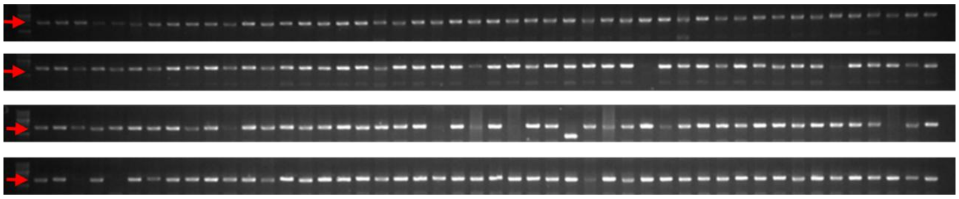

**B**

| Index | Combination | N-linker_sequenced | C-linker_sequenced |
|---|---|---|---|
| 1 | N16C4 | GGSGGSGGGGSGGGGS | SGGG |
| 2 | N16C7 | GGSGGSGGGGSGGGGS | SGGGSGG |
| 3 | N3C14 | GGS | SGGGGSGGGGSGGG |
| 4 | N4C20 | GGGS | SGGGGSGGGGSGGGGSGGGG |
| 5 | N16C19 | Frame shift | Frame shift |
| 6 | N9C19 | GGGSGGGGS | SGGGGSGGGGSGGGGSGGG |
| 7 | N17C17 | GGSGGGSGGGGSGGGGS | SGGGGSGGGGSGGGSGG |
| 8 | N20C20 | GGGGSGGGGSGGGGSGGGGS | SGGGGSGGGGSGGGGSGGGG |
| 9 | N5C12 | GGGGS | SGGGGSGGGSGG |
| 10 | N5C13 | GGGGS | SGGGGSGGGGSGG |
| 11 | N6C10 | GGSGGS | SGGGGSGGGG |
| 12 | N8C7 | GGSGGGGS | SGGGSGG |
| 13 | N9C7 | GGGSGGGGS | SGGGSGG |
| 14 | N11C12 | GGSGGSGGGGS | SGGGGSGGGSGG |
| 15 | N12C5 | GGSGGGSGGGGS | SGGGG |
| 16 | N13C9 | GGSGGGGSGGGGS | SGGGGSGGG |
| 17 | N18C18 | GGSGGGGSGGGGSGGGGS | SGGGGSGGGGSGGGGSGG |
| 18 | N18C19 | GGSGGGGSGGGGSGGGGS | SGGGGSGGGGSGGGGSGGG |
| 19 | N18C20 | GGSGGGGSGGGGSGGGGS | SGGGGSGGGGSGGGGSGGGG |
| 20 | N19C18 | GGGSGGGGSGGGGSGGGGS | SGGGGSGGGGSGGGGSGG |
| 21 | N19C19 | GGGSGGGGSGGGGSGGGGS | SGGGGSGGGGSGGGGSGGG |
| 22 | N19C20 | GGGSGGGGSGGGGSGGGGS | SGGGGSGGGGSGGGGSGGGG |
| 23 | N20C18 | GGGGSGGGGSGGGGSGGGGS | SGGGGSGGGGSGGGGSGG |
| 24 | N20C19 | GGGGSGGGGSGGGGSGGGGS | SGGGGSGGGGSGGGGSGGG |

**Supplementary Fig. 1.** Cloning efficiency analysis. A, Agarose gel image of colony PCR of randomly picked 196 clones after ligation (96 randomly picked variants x 4 clones per variant). Red arrows point to the band of interest (~1600 bp). B, Sanger sequencing results of 24 variants; 1 clone per variant.

>CA14-stabilized
QVQLQESGGGLVQPGGSLRLSCAASGSTSRQYDMGWFRQAPGKEREFVSAISSNQDQPPYYA
DSVKGRFTISRDNAKNTVYLQMNSLKPEDTATYYCAFKQHHANGAYWGQGTQVTVSS
>DB21-stabilized
QVQLQESGGGLVQPGGSLRLSCAASGTTYGQTNMGWFRQAPGKEREFVSAISGLQGRDLYYAD
SVKGRFTISRDNAKNTVYLQMNSLKPEDTATYYCAFHDFLRMWEYWGQGTQVTVSS

**Supplementary Fig. 2.** Stabilized nanobody amino acid sequences

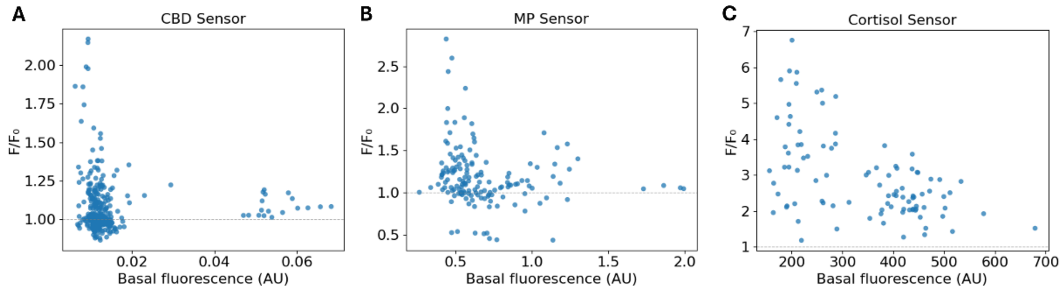


**Supplementary Fig. 3.** Correlation between basal fluorescence and $\Delta F/F_0$ across linker-length variants. A, the scatter plot for CBD sensor variants. B, the scatter plot for MP sensor variants. C, the scatter plot for cortisol sensor variants.

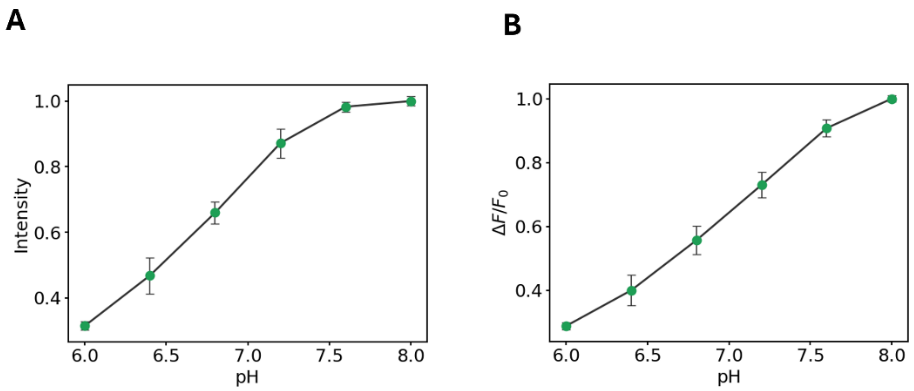


**Supplementary Fig.4.** Biophysical characterization of N12C2 CBD sensor variant. A, Fluorescence intensity of the N12C2 sensor measured across a range of pH values. B, Ligand-dependent fluorescence response of the N12C2 sensor as a function of pH. Experiments were independently repeated three times with similar results.

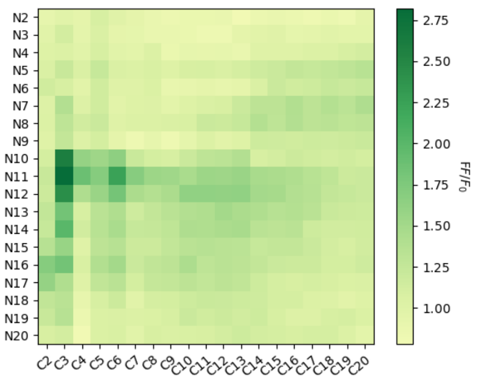


**Supplementary Fig. 5.** Exhaustive evaluation of the MP linker-length matrix. Heatmap of ligand-dependent fluorescence response of all 361 linker variants of the MP sensor, expressed as $F/F_0$ (fluorescence at saturating ligand divided by basal fluorescence).

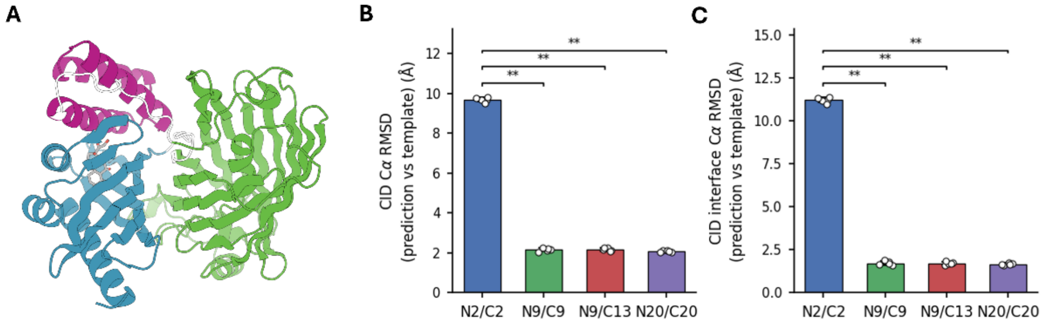


**Supplementary Fig. 6.** Structure prediction of cortisol N9C13 variant. A, Representative predicted structure of N9C13 variant. Green: cpGFP; cyan: mhcy129; magenta: CorD1. B, CID Cα RMSD between the predicted sensor structure and the predicted CID structure for four variants of linker combinations: N2C2, N9C9, N9C13 and N20C20. C, CID interface Cα RMSD between the predicted sensor structure and the predicted CID structure for four variants of linker combinations: N2C2, N9C9, N9C13 and N20C20. cpGFP crystal structure (PDB: 3WLC) was used as a constraint for Chai-1 model using ESM embedding. 5 structures were generated per variant. The statistics is using pairwise Mann-Whitney U tests across all 6 linker pairs per panel. Only significant pairs (* for $p < 0.05$; ** for $p < 0.01$; *** for $p < 0.001$) are drawn.Fig. S2. Stabilized nanobody amino acid sequences

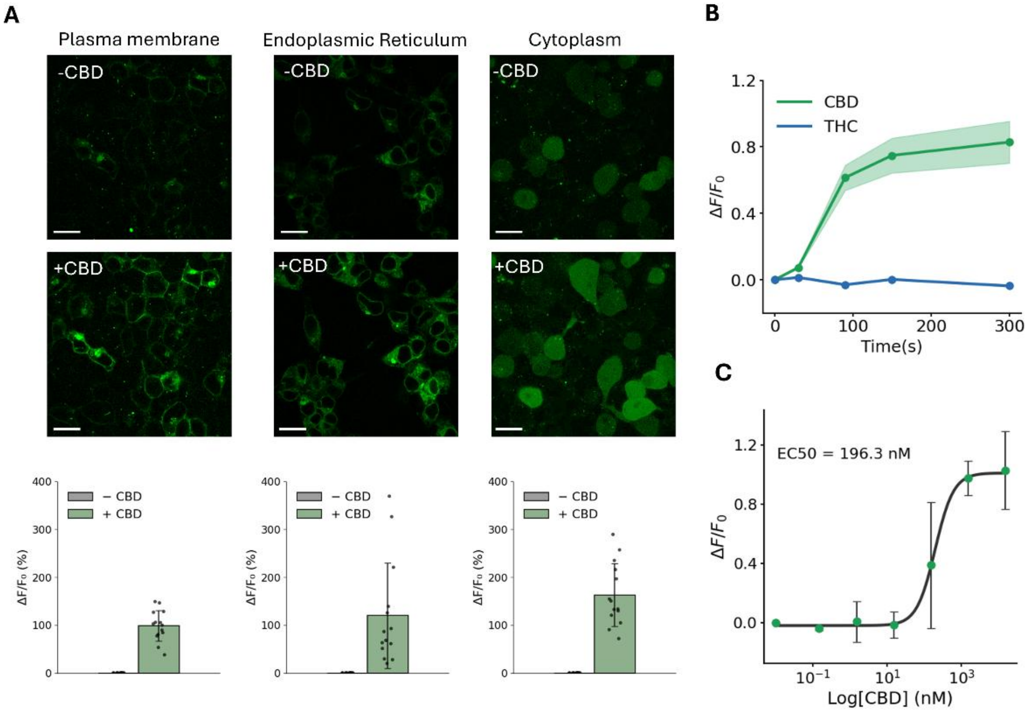


**Supplementary Fig. 7.** Subcellular localization and fluorescence response of the CBD N12C2 sensor. A, Representative fluorescence images of cells expressing the CBD N12C2 sensor in the plasma membrane (left), ER lumen (middle), and cytosol (right) before and after application of 30 μM CBD. Bottom panels show $\Delta F/F_0$ of individual cells (ROIs), with bars representing mean ± s.d. Individual data points represent single-cell measurements. Scale bar= 40 μm. B, Quantification of $\Delta F/F_0$ over time following addition of 30 μM CBD or THC. Data represent mean ± s.d. of 14 cells shown in Figure. 7B, tracked across all time points. C, Representative dose–response curve measured from the mean fluorescence intensity of threshold-based masks encompassing all cells in a field of view, across a range of CBD concentrations. Data represent mean ± s.d. from three independent biological replicates (independent transfections). $EC_{50}$ = 197 nM; maximum $\Delta F/F_0$ = 103%.

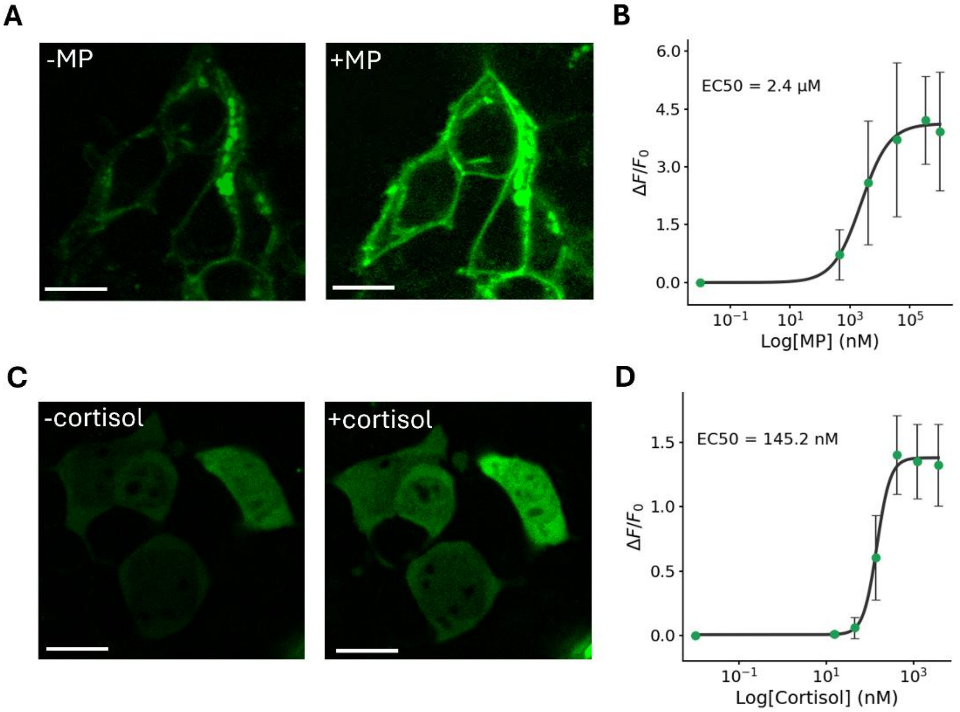


**Supplementary Fig. 8.** Cellular fluorescence response of the MP N11C3 sensor and the cortisol N5C8 sensor.
A, Representative fluorescence images showing MP sensor N11C3 expressed on the plasma membrane before ligand addition (left), after addition of 1 mM MP (right). Scale bar, 20 μm.
B, Dose–response curve of the MP sensor N11C3 measured from the mean fluorescence intensity of threshold-based masks encompassing all cells in a field of view, across a range of concentrations of ligands. $EC_{50}$ = 2.4 μM, maximum ΔF/F0 = 420%. Data represent mean ± s.d. of three biological replicates.
C, Representative fluorescence images showing cortisol sensor N5C8 expressed cytosolically before ligand addition (left), after addition of 100 nM cortisol (right). Scale bar, 20 μm.
D, Dose–response curve of the cortisol sensor N5C8 measured from the mean fluorescence intensity of threshold-based masks encompassing all cells in a field of view, across a range of concentrations of ligands. $EC_{50}$ = 145.2 nM, maximum $\Delta F/F_0$ = 140%. Data represent mean ± s.d. of three biological replicates.

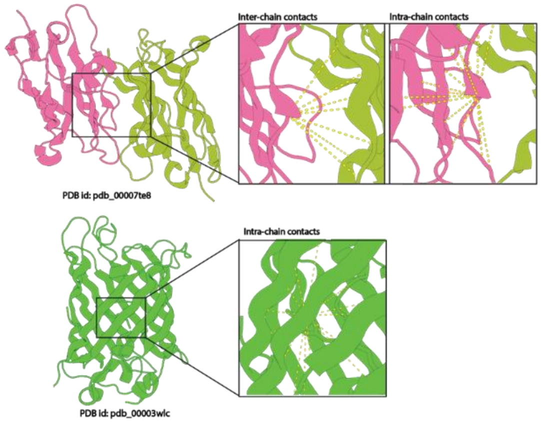


**Supplementary Fig. 9.** Structural contacts used as constraints for Chai1-based structural prediction. Within the CID complex (PDB: 7TE8), both intra- and inter-chain residue (Cα) contacts within 11 A were used; within the cpGFP (PDB: 3WLC), intra-chain residue (Cα) contacts within 11 A were used.

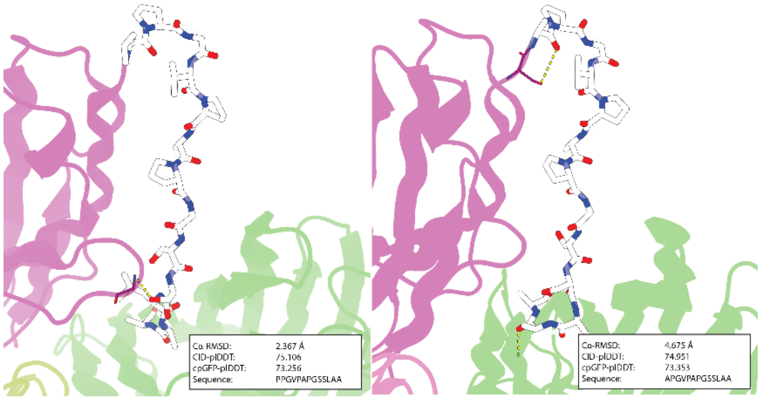


**Supplementary Fig. 10.** Predicted structure and filtering metrics of the top two linker designs. Structural confidence and conformational metrics (Cα-RMSD, CID-pLDDT, and cpGFP-pLDDT) together with linker sequences for the top two designed variants (LDc1, left; LDc2, right).

| Name | DNA Sequence |
|---|---|
| VgsVLR-R2 | caccctgacctacggcgtgcagtgcttcagccgctaccccgaccacatgaagcagcacgacttcttcaagtccgccatgcccgaaggctacattcaggagcgcaccatcttcttcaaggacgacggcaactataagacacgcgctgaggttaagttcgagggcgacactctggttaaccgcatcgagctgaagggcatcgacttcaaggaggacggcaacatcctgggccataagcttgaatataacttcaacggcggtttGTCTTCCAAACGAGGTGATTTCAACAAATTTTGTTGATGAT |
| VgsVLR-R3 | caccctgacctacggcgtgcagtgcttcagccgctaccccgaccacatgaagcagcacgacttcttcaagtccgccatgcccgaaggctacattcaggagcgcaccatcttcttcaaggacgacggcaactataagacacgcgctgaggttaagttcgagggcgacactctggttaaccgcatcgagctgaagggcatcgacttcaaggaggacggcaacatcctgggccataagcttgaatataacttcaacTCCggcggtttGTCTTCCAAACGAGGTGATTTCAACAAATTTTGTTGATGAT |
| VgsVLR-R4 | caccctgacctacggcgtgcagtgcttcagccgctaccccgaccacatgaagcagcacgacttcttcaagtccgccatgcccgaaggctacattcaggagcgcaccatcttcttcaaggacgacggcaactataagacacgcgctgaggttaagttcgagggcgacactctggttaaccgcatcgagctgaagggcatcgacttcaaggaggacggcaacatcctgggccataagcttgaatataacttcaacTCCGGAggcggtttGTCTTCCAAACGAGGTGATTTCAACAAATTTTGTTGATGAT |
| VgsVLR-R5 | caccctgacctacggcgtgcagtgcttcagccgctaccccgaccacatgaagcagcacgacttcttcaagtccgccatgcccgaaggctacattcaggagcgcaccatcttcttcaaggacgacggcaactataagacacgcgctgaggttaagttcgagggcgacactctggttaaccgcatcgagctgaagggcatcgacttcaaggaggacggcaacatcctgggccataagcttgaatataacttcaacTCCGGAGGTggcggtttGTCTTCCAAACGAGGTGATTTCAACAAATTTTGTTGATGAT |
| VgsVLR-R6 | caccctgacctacggcgtgcagtgcttcagccgctaccccgaccacatgaagcagcacgacttcttcaagtccgccatgcccgaaggctacattcaggagcgcaccatcttcttcaaggacgacggcaactataagacacgcgctgaggttaagttcgagggcgacactctggttaaccgcatcgagctgaagggcatcgacttcaaggaggacggcaacatcctgggccataagcttgaatataacttcaacTCCGGAGGTTCTggcggtttGTCTTCCAAACGAGGTGATTTCAACAAATTTTGTTGATGAT |
| VgsVLR-R7 | caccctgacctacggcgtgcagtgcttcagccgctaccccgaccacatgaagcagcacgacttcttcaagtccgccatgcccgaaggctacattcaggagcgcaccatcttcttcaaggacgacggcaactataagacacgcgctgaggttaagttcgagggcgacactctggttaaccgcatcgagctgaagggcatcgacttcaaggaggacggcaacatcctgggccataagcttgaatataacttcaacTCCGGAGGTGGCTCTggcggtttGTCTTCCAAACGAGGTGATTTCAACAAATTTTGTTGATGAT |
| VgsVLR-R8 | caccctgacctacggcgtgcagtgcttcagccgctaccccgaccacatgaagcagcacgacttcttcaagtccgccatgcccgaaggctacattcaggagcgcaccatcttcttcaaggacgacggcaactataagacacgcgctgaggttaagttcgagggcgacactctggttaaccgcatcgagctgaagggcatcgacttcaaggaggacggcaacatcctgggccataagcttgaatataacttcaacTCCGGAGGTGGCGGATCTggcggtttGTCTTCCAAACGAGGTGATTTCAACAAATTTTGTTGATGAT |
| VgsVLR-R9 | caccctgacctacggcgtgcagtgcttcagccgctaccccgaccacatgaagcagcacgacttcttcaagtccgccatgcccgaaggctacattcaggagcgcaccatcttcttcaaggacgacggcaactataagacacgcgctgaggttaagttcgagggcgacactctggttaaccgcatcgagctgaagggcatcgacttcaaggaggacggcaacatcctgggccataagcttgaatataacttcaacTCCGGAGGTGGCGGATCTGGTggcggtttGTCTTCCAAACGAGGTGATTTCAACAAATTTTGTTGATGAT |
| VgsVLR-R10 | caccctgacctacggcgtgcagtgcttcagccgctaccccgaccacatgaagcagcacgacttcttcaagtccgccatgcccgaaggctacattcaggagcgcaccatcttcttcaaggacgacggcaactataagacacgcgctgaggttaagttcgagggcgacactctggttaaccgcatcgagctgaagggcatcgacttcaaggaggacggcaacatcctgggccataagcttgaatataacttcaacTCCGGAGGTGGCGGATCTGGTGGAggcggtttGTCTTCCAAACGAGGTGATTTCAACAAATTTTGTTGATGAT |
| VgsVLR-R11 | caccctgacctacggcgtgcagtgcttcagccgctaccccgaccacatgaagcagcacgacttcttcaagtccgccatgcccgaaggctacattcaggagcgcaccatcttcttcaaggacgacggcaactataagacacgcgctgaggttaagttcgagggcgacactctggttaaccgcatcgagctgaagggcatcgacttcaaggaggacggcaacatcctgggccataagcttgaatataacttcaacTCCGGAGGTGGCGGATCTGGTGGAAGTggcggtttGTCTTCCAAACGAGGTGATTTCAACAAATTTTGTTGATGAT |

| | |
|---|---|
| VgsVLR-R12 | caccctgacctacggcgtgcagtgcttcagccgctaccccgaccacatgaagcagcacgacttcttcaagtccgccatgcccgaaggctacattcaggagcgcaccatcttcttcaaggacgacggcaactataagacacgcgctgaggttaagttcgagggcgacactctggttaaccgcatcgagctgaagggcatcgacttcaaggaggacggcaacatcctgggccataagcttgaatataacttcaacTCCGGAGGTGGCGGATCTGGTGGAGGAAGTggcggtttGTCTTCCAAACGAGGTGATTTCAACAAATTTTGTTGATGAT |
| VgsVLR-R13 | caccctgacctacggcgtgcagtgcttcagccgctaccccgaccacatgaagcagcacgacttcttcaagtccgccatgcccgaaggctacattcaggagcgcaccatcttcttcaaggacgacggcaactataagacacgcgctgaggttaagttcgagggcgacactctggttaaccgcatcgagctgaagggcatcgacttcaaggaggacggcaacatcctgggccataagcttgaatataacttcaacTCCGGAGGTGGCGGATCTGGTGGAGGAGGCAGTggcggtttGTCTTCCAAACGAGGTGATTTCAACAAATTTTGTTGATGAT |
| VgsVLR-R14 | caccctgacctacggcgtgcagtgcttcagccgctaccccgaccacatgaagcagcacgacttcttcaagtccgccatgcccgaaggctacattcaggagcgcaccatcttcttcaaggacgacggcaactataagacacgcgctgaggttaagttcgagggcgacactctggttaaccgcatcgagctgaagggcatcgacttcaaggaggacggcaacatcctgggccataagcttgaatataacttcaacTCCGGAGGTGGCGGATCTGGTGGAGGAGGCAGTGGCggcggtttGTCTTCCAAACGAGGTGATTTCAACAAATTTTGTTGATGAT |
| VgsVLR-R15 | caccctgacctacggcgtgcagtgcttcagccgctaccccgaccacatgaagcagcacgacttcttcaagtccgccatgcccgaaggctacattcaggagcgcaccatcttcttcaaggacgacggcaactataagacacgcgctgaggttaagttcgagggcgacactctggttaaccgcatcgagctgaagggcatcgacttcaaggaggacggcaacatcctgggccataagcttgaatataacttcaacTCCGGAGGTGGCGGATCTGGTGGAGGAGGCAGTGGCGGTggcggtttGTCTTCCAAACGAGGTGATTTCAACAAATTTTGTTGATGAT |
| VgsVLR-R16 | caccctgacctacggcgtgcagtgcttcagccgctaccccgaccacatgaagcagcacgacttcttcaagtccgccatgcccgaaggctacattcaggagcgcaccatcttcttcaaggacgacggcaactataagacacgcgctgaggttaagttcgagggcgacactctggttaaccgcatcgagctgaagggcatcgacttcaaggaggacggcaacatcctgggccataagcttgaatataacttcaacTCCGGAGGTGGCGGATCTGGTGGAGGAGGCAGTGGCGGTAGCggcggtttGTCTTCCAAACGAGGTGATTTCAACAAATTTTGTTGATGAT |
| VgsVLR-R17 | caccctgacctacggcgtgcagtgcttcagccgctaccccgaccacatgaagcagcacgacttcttcaagtccgccatgcccgaaggctacattcaggagcgcaccatcttcttcaaggacgacggcaactataagacacgcgctgaggttaagttcgagggcgacactctggttaaccgcatcgagctgaagggcatcgacttcaaggaggacggcaacatcctgggccataagcttgaatataacttcaacTCCGGAGGTGGCGGATCTGGTGGAGGAGGCAGTGGCGGTGGAAGCggcggtttGTCTTCCAAACGAGGTGATTTCAACAAATTTTGTTGATGAT |
| VgsVLR-R18 | caccctgacctacggcgtgcagtgcttcagccgctaccccgaccacatgaagcagcacgacttcttcaagtccgccatgcccgaaggctacattcaggagcgcaccatcttcttcaaggacgacggcaactataagacacgcgctgaggttaagttcgagggcgacactctggttaaccgcatcgagctgaagggcatcgacttcaaggaggacggcaacatcctgggccataagcttgaatataacttcaacTCCGGAGGTGGCGGATCTGGTGGAGGAGGCAGTGGCGGTGGAGGCAGCggcggtttGTCTTCCAAACGAGGTGATTTCAACAAATTTTGTTGATGAT |
| VgsVLR-R19 | caccctgacctacggcgtgcagtgcttcagccgctaccccgaccacatgaagcagcacgacttcttcaagtccgccatgcccgaaggctacattcaggagcgcaccatcttcttcaaggacgacggcaactataagacacgcgctgaggttaagttcgagggcgacactctggttaaccgcatcgagctgaagggcatcgacttcaaggaggacggcaacatcctgggccataagcttgaatataacttcaacTCCGGAGGTGGCGGATCTGGTGGAGGAGGCAGTGGCGGTGGAGGCAGCGGAggcggtttGTCTTCCAAACGAGGTGATTTCAACAAATTTTGTTGATGAT |
| VgsVLR-R20 | caccctgacctacggcgtgcagtgcttcagccgctaccccgaccacatgaagcagcacgacttcttcaagtccgccatgcccgaaggctacattcaggagcgcaccatcttcttcaaggacgacggcaactataagacacgcgctgaggttaagttcgagggcgacactctggttaaccgcatcgagctgaagggcatcgacttcaaggaggacggcaacatcctgggccataagcttgaatataacttcaacTCCGGAGGTGGCGGATCTGGTGGAGGAGGCAGTGGCGGTGGAGGCAGCGGAGGCggcggtttGTCTTCCAAACGAGGTGATTTCAACAAATTTTGTTGATGAT |

| | |
|---|---|
| VgsVLR-F2 | TCAGTAATTTTCCAATATGAGAACATCAATTGTACGAAGACggtggctataacgtctttatcatggccgacaagcagaagaacggcatcaaggcgaacttcaagatccgccacaacatcgaggacggcggcgtgcagctcgcctatcactaccagcagaacacccccatcggcgacggccccgtgctgctgcccgacaaccactacctgagcgtgcagtccaaactgagcaaagaccccaacgagaagcgcgatcacatggtcctgctggagttcgtgaccgccgccgggatcactctcg |
| VgsVLR-F3 | TCAGTAATTTTCCAATATGAGAACATCAATTGTACGAAGACggtggcAGTtataacgtctttatcatggccgacaagcagaagaacggcatcaaggcgaacttcaagatccgccacaacatcgaggacggcggcgtgcagctcgcctatcactaccagcagaacacccccatcggcgacggccccgtgctgctgcccgacaaccactacctgagcgtgcagtccaaactgagcaaagaccccaacgagaagcgcgatcacatggtcctgctggagttcgtgaccgccgccgggatcactctcg |
| VgsVLR-F4 | TCAGTAATTTTCCAATATGAGAACATCAATTGTACGAAGACggtggcGGAAGTtataacgtctttatcatggccgacaagcagaagaacggcatcaaggcgaacttcaagatccgccacaacatcgaggacggcggcgtgcagctcgcctatcactaccagcagaacacccccatcggcgacggccccgtgctgctgcccgacaaccactacctgagcgtgcagtccaaactgagcaaagaccccaacgagaagcgcgatcacatggtcctgctggagttcgtgaccgccgccgggatcactctcg |
| VgsVLR-F5 | TCAGTAATTTTCCAATATGAGAACATCAATTGTACGAAGACggtggcGGCGGAAGTtataacgtctttatcatggccgacaagcagaagaacggcatcaaggcgaacttcaagatccgccacaacatcgaggacggcggcgtgcagctcgcctatcactaccagcagaacacccccatcggcgacggccccgtgctgctgcccgacaaccactacctgagcgtgcagtccaaactgagcaaagaccccaacgagaagcgcgatcacatggtcctgctggagttcgtgaccgccgccgggatcactctcg |
| VgsVLR-F6 | TCAGTAATTTTCCAATATGAGAACATCAATTGTACGAAGACggtggcTCCGGCGGAAGTtataacgtctttatcatggccgacaagcagaagaacggcatcaaggcgaacttcaagatccgccacaacatcgaggacggcggcgtgcagctcgcctatcactaccagcagaacacccccatcggcgacggccccgtgctgctgcccgacaaccactacctgagcgtgcagtccaaactgagcaaagaccccaacgagaagcgcgatcacatggtcctgctggagttcgtgaccgccgccgggatcactctcg |
| VgsVLR-F7 | TCAGTAATTTTCCAATATGAGAACATCAATTGTACGAAGACggtggcTCCGGAGGCGGAAGTtataacgtctttatcatggccgacaagcagaagaacggcatcaaggcgaacttcaagatccgccacaacatcgaggacggcggcgtgcagctcgcctatcactaccagcagaacacccccatcggcgacggccccgtgctgctgcccgacaaccactacctgagcgtgcagtccaaactgagcaaagaccccaacgagaagcgcgatcacatggtcctgctggagttcgtgaccgccgccgggatcactctcg |
| VgsVLR-F8 | TCAGTAATTTTCCAATATGAGAACATCAATTGTACGAAGACggtggcTCCGGTGGAGGCGGAAGTtataacgtctttatcatggccgacaagcagaagaacggcatcaaggcgaacttcaagatccgccacaacatcgaggacggcggcgtgcagctcgcctatcactaccagcagaacacccccatcggcgacggccccgtgctgctgcccgacaaccactacctgagcgtgcagtccaaactgagcaaagaccccaacgagaagcgcgatcacatggtcctgctggagttcgtgaccgccgccgggatcactctcg |
| VgsVLR-F9 | TCAGTAATTTTCCAATATGAGAACATCAATTGTACGAAGACggtggcGGCTCCGGTGGAGGCGGAAGTtataacgtctttatcatggccgacaagcagaagaacggcatcaaggcgaacttcaagatccgccacaacatcgaggacggcggcgtgcagctcgcctatcactaccagcagaacacccccatcggcgacggccccgtgctgctgcccgacaaccactacctgagcgtgcagtccaaactgagcaaagaccccaacgagaagcgcgatcacatggtcctgctggagttcgtgaccgccgccgggatcactctcg |
| VgsVLR-F10 | TCAGTAATTTTCCAATATGAGAACATCAATTGTACGAAGACggtggcGGTGGCTCCGGTGGAGGCGGAAGTtataacgtctttatcatggccgacaagcagaagaacggcatcaaggcgaacttcaagatccgccacaacatcgaggacggcggcgtgcagctcgcctatcactaccagcagaacacccccatcggcgacggccccgtgctgctgcccgacaaccactacctgagcgtgcagtccaaactgagcaaagaccccaacgagaagcgcgatcacatggtcctgctggagttcgtgaccgccgccgggatcactctcg |
| VgsVLR-F11 | TCAGTAATTTTCCAATATGAGAACATCAATTGTACGAAGACggtggcTCTGGTGGCTCCGGTGGAGGCGGAAGTtataacgtctttatcatggccgacaagcagaagaacggcatcaaggcgaacttcaagatccgccacaacatcgaggacggcggcgtgcagctcgcctatcactaccagcagaacacccccatcggcgacggccccgtgctgctgcccgacaaccactacctgagcgtgcagtccaaactgagcaaagaccccaacgagaagcgcgatcacatggtcctgctggagttcgtgaccgccgccgggatcactctcg |

| | |
|---|---|
| VgsVLR<br>-F12 | TCAGTAATTTTCCAATATGAGAACATCAATTGTACGAAGACggtggcTCTGGAGGTGGCTCC<br>GGTGGAGGCGGAAGTtataacgtctttatcatggccgacaagcagaagaacggcatcaaggcgaacttcaagatccg<br>ccacaacatcgaggacggcggcgtgcagctcgcctatcactaccagcagaacacccccatcggcgacggccccgtgctgctgc<br>ccgacaaccactacctgagcgtgcagtccaaactgagcaaagaccccaacgagaagcgcgatcacatggtcctgctggagttc<br>gtgaccgccgccgggatcactctcg |
| VgsVLR<br>-F13 | TCAGTAATTTTCCAATATGAGAACATCAATTGTACGAAGACggtggcTCTGGAGGAGGTGG<br>CTCCGGTGGAGGCGGAAGTtataacgtctttatcatggccgacaagcagaagaacggcatcaaggcgaacttcaa<br>gatccgccacaacatcgaggacggcggcgtgcagctcgcctatcactaccagcagaacacccccatcggcgacggccccgtg<br>ctgctgcccgacaaccactacctgagcgtgcagtccaaactgagcaaagaccccaacgagaagcgcgatcacatggtcctgct<br>ggagttcgtgaccgccgccgggatcactctcg |
| VgsVLR<br>-F14 | TCAGTAATTTTCCAATATGAGAACATCAATTGTACGAAGACggtggcGGGTCTGGAGGAGG<br>TGGCTCCGGTGGAGGCGGAAGTtataacgtctttatcatggccgacaagcagaagaacggcatcaaggcgaact<br>tcaagatccgccacaacatcgaggacggcggcgtgcagctcgcctatcactaccagcagaacacccccatcggcgacggccc<br>cgtgctgctgcccgacaaccactacctgagcgtgcagtccaaactgagcaaagaccccaacgagaagcgcgatcacatggtcc<br>tgctggagttcgtgaccgccgccgggatcactctcg |
| VgsVLR<br>-F15 | TCAGTAATTTTCCAATATGAGAACATCAATTGTACGAAGACggtggcGGAGGGTCTGGAGG<br>AGGTGGCTCCGGTGGAGGCGGAAGTtataacgtctttatcatggccgacaagcagaagaacggcatcaaggc<br>gaacttcaagatccgccacaacatcgaggacggcggcgtgcagctcgcctatcactaccagcagaacacccccatcggcgac<br>ggccccgtgctgctgcccgacaaccactacctgagcgtgcagtccaaactgagcaaagaccccaacgagaagcgcgatcaca<br>tggtcctgctggagttcgtgaccgccgccgggatcactctcg |
| VgsVLR<br>-F16 | TCAGTAATTTTCCAATATGAGAACATCAATTGTACGAAGACggtggcAGCGGAGGGTCTGG<br>AGGAGGTGGCTCCGGTGGAGGCGGAAGTtataacgtctttatcatggccgacaagcagaagaacggcatca<br>aggcgaacttcaagatccgccacaacatcgaggacggcggcgtgcagctcgcctatcactaccagcagaacacccccatcgg<br>cgacggccccgtgctgctgcccgacaaccactacctgagcgtgcagtccaaactgagcaaagaccccaacgagaagcgcgat<br>cacatggtcctgctggagttcgtgaccgccgccgggatcactctcg |
| VgsVLR<br>-F17 | TCAGTAATTTTCCAATATGAGAACATCAATTGTACGAAGACggtggcAGCGGTGGAGGGTC<br>TGGAGGAGGTGGCTCCGGTGGAGGCGGAAGTtataacgtctttatcatggccgacaagcagaagaacggc<br>atcaaggcgaacttcaagatccgccacaacatcgaggacggcggcgtgcagctcgcctatcactaccagcagaacacccccat<br>cggcgacggccccgtgctgctgcccgacaaccactacctgagcgtgcagtccaaactgagcaaagaccccaacgagaagcg<br>cgatcacatggtcctgctggagttcgtgaccgccgccgggatcactctcg |
| VgsVLR<br>-F18 | TCAGTAATTTTCCAATATGAGAACATCAATTGTACGAAGACggtggcAGCGGCGGTGGAGG<br>GTCTGGAGGAGGTGGCTCCGGTGGAGGCGGAAGTtataacgtctttatcatggccgacaagcagaagaa<br>cggcatcaaggcgaacttcaagatccgccacaacatcgaggacggcggcgtgcagctcgcctatcactaccagcagaacacc<br>cccatcggcgacggccccgtgctgctgcccgacaaccactacctgagcgtgcagtccaaactgagcaaagaccccaacgaga<br>agcgcgatcacatggtcctgctggagttcgtgaccgccgccgggatcactctcg |
| VgsVLR<br>-F19 | TCAGTAATTTTCCAATATGAGAACATCAATTGTACGAAGACggtggcGGTAGCGGCGGTGG<br>AGGGTCTGGAGGAGGTGGCTCCGGTGGAGGCGGAAGTtataacgtctttatcatggccgacaagcaga<br>agaacggcatcaaggcgaacttcaagatccgccacaacatcgaggacggcggcgtgcagctcgcctatcactaccagcagaa<br>cacccccatcggcgacggccccgtgctgctgcccgacaaccactacctgagcgtgcagtccaaactgagcaaagaccccaac<br>gagaagcgcgatcacatggtcctgctggagttcgtgaccgccgccgggatcactctcg |
| VgsVLR<br>-F20 | TCAGTAATTTTCCAATATGAGAACATCAATTGTACGAAGACggtggcGGAGGTAGCGGCGG<br>TGGAGGGTCTGGAGGAGGTGGCTCCGGTGGAGGCGGAAGTtataacgtctttatcatggccgacaag<br>cagaagaacggcatcaaggcgaacttcaagatccgccacaacatcgaggacggcggcgtgcagctcgcctatcactaccagc<br>agaacacccccatcggcgacggccccgtgctgctgcccgacaaccactacctgagcgtgcagtccaaactgagcaaagaccc<br>caacgagaagcgcgatcacatggtcctgctggagttcgtgaccgccgccgggatcactctcg |

| | |
|---|---|
| VC155-M-Ext | cactgcacgccgtaggtcagggtggtcacgagggtgggccagggcacgggcagcttgccggtggtgcagatgaacttcagggtcagcttgccgtaggtggcatcgccctcgccctcgccggacacgctgaacttgtggccgtttacgtcgccgtccagctcgaccaggatgggcaccaccccggtgaacagctcctcgcccttgctcaccatgctccctccggtaccgcccttgtacagctcgtccatgccgagagtgatcccggcggcgg |

**Supplementary Table 1.** All starting DNA fragments used to generate the cpGFP-linker length library. Each overlap extension PCR reaction contains a left fragment (VgsVL-F, corresponding to N-linkers), a right fragment (VgsVL-R, corresponding to C-linkers) and a middle fragment (VC155-M-Ext).

| Construct | Amino Acid Sequence |
|---|---|
| >CBD-N12C2 | MQVQLQESGGGLVQPGGSLRLSCAASGSTSRQYDMGWFRQAPGKEREFVSAISSNQDQPPYYADSVKGRFTISRDNAKNTVYLQMNSLKPEDTATYYCAFKQHHANGAYWGQGTQVTVSSGGSGGGSGGGGSYNVFIMADKQKNGIKANFKIRHNIEDGGVQLAYHYQQNTPIGDGPVLLPDNHYLSVQSKLSKDPNEKRDHMVLLEFVTAAGITLGMDELYKGGTGGSMVSKGEELFTGVVPILVELDGDVNGHKFSVSGEGEGDATYGKLTLKFICTTGKLPVPWPTLVTTLTYGVQCFSRYPDHMKQHDFFKSAMPEGYIQERTIFFKDDGNYKTRAEVKFEGDTLVNRIELKGIDFKEDGNILGHKLEYNFNGGQVQLQESGGGLVQPGGSLRLSCAASGTTYGQTNMGWFRQAPGKEREFVSAISGLQGRDLYYADSVKGRFTISRDNAKNTVYLQMNSLKPEDTATYYCAFHDFLRMWEYWGQGTQVTVSS |
| >Cortisol-N5C8 | MTSSKEAEEAIRDMLRRWYEAINKGDMEKLKSLVDPDASFHFAITNQQYDKEQFLEMIKEALKQDLKVEVKSIHIQQQPRGDHVTVTVHVEAHMNQNGQTHTFTVTDHYHFVRKGDSWKITRTQWHIHLQGGGGSYNVFIMADKQKNGIKANFKIRHNIEDGGVQLAYHYQQNTPIGDGPVLLPDNHYLSVQSKLSKDPNEKRDHMVLLEFVTAAGITLGMDELYKGGTGGSMVSKGEELFTGVVPILVELDGDVNGHKFSVSGEGEGDATYGKLTLKFICTTGKLPVPWPTLVTTLTYGVQCFSRYPDHMKQHDFFKSAMPEGYIQERTIFFKDDGNYKTRAEVKFEGDTLVNRIELKGIDFKEDGNILGHKLEYNFNSGGGGSGGMDKSKLTLKVFNIAMEYGLNDEQYQQLADFAFKKLEEGKSLEEIEKELREKAKELA |
| >CBD-N12C2-LDc1 | MQVQLQESGGGLVQPGGSLRLSCAASGSTSRQYDMGWFRQAPGKEREFVSAISSNQDQPPYYADSVKGRFTISRDNAKNTVYLQMNSLKPEDTATYYCAFKQHHANGAYWGQGTQVTVSPPGVPAPGSSLAAYNVFIMADKQKNGIKANFKIRHNIEDGGVQLAYHYQQNTPIGDGPVLLPDNHYLSVQSKLSKDPNEKRDHMVLLEFVTAAGITLGMDELYKGGTGGSMVSKGEELFTGVVPILVELDGDVNGHKFSVSGEGEGDATYGKLTLKFICTTGKLPVPWPTLVTTLTYGVQCFSRYPDHMKQHDFFKSAMPEGYIQERTIFFKDDGNYKTRAEVKFEGDTLVNRIELKGIDFKEDGNILGHKLEYNFNGGQVQLQESGGGLVQPGGSLRLSCAASGTTYGQTNMGWFRQAPGKEREFVSAISGLQGRDLYYADSVKGRFTISRDNAKNTVYLQMNSLKPEDTATYYCAFHDFLRMWEYWGQGTQVTVSS |
| >CBD-N12C2-LDc2 | MQVQLQESGGGLVQPGGSLRLSCAASGSTSRQYDMGWFRQAPGKEREFVSAISSNQDQPPYYADSVKGRFTISRDNAKNTVYLQMNSLKPEDTATYYCAFKQHHANGAYWGQGTQVTVSAPGVPAPGSSLAAYNVFIMADKQKNGIKANFKIRHNIEDGGVQLAYHYQQNTPIGDGPVLLPDNHYLSVQSKLSKDPNEKRDHMVLLEFVTAAGITLGMDELYKGGTGGSMVSKGEELFTGVVPILVELDGDVNGHKFSVSGEGEGDATYGKLTLKFICTTGKLPVPWPTLVTTLTYGVQCFSRYPDHMKQHDFFKSAMPEGYIQERTIFFKDDGNYKTRAEVKFEGDTLVNRIELKGIDFKEDGNILGHKLEYNFNGGQVQLQESGGGLVQPGGSLRLSCAASGTTYGQTNMGWFRQAPGKEREFVSAISGLQGRDLYYADSVKGRFTISRDNAKNTVYLQMNSLKPEDTATYYCAFHDFLRMWEYWGQGTQVTVSS |

**Supplementary Table 2.** Amino acid sequences of the sensor variants
